# The ORB2 RNA-binding protein negatively regulates its target transcripts during the Drosophila maternal-to-zygotic transition via its functionally conserved Zinc-binding ‘ZZ’ domain

**DOI:** 10.1101/2025.07.10.664187

**Authors:** Timothy C.H. Low, Hua Luo, Chalini Weerasooriya, Zhuyi Wang, Sichun Lin, Angelo Karaiskakis, Stephane Angers, Craig A. Smibert, Howard D. Lipshitz

## Abstract

Post-transcriptional regulation is particularly prominent during the maternal-to-zygotic transition (MZT), a developmental phase during which a large proportion of maternally provided mRNAs is repressed and cleared from metazoan embryos. RNA-binding proteins (RBPs) are key components of the post-transcriptional regulatory machinery. We show that the ORB2 RBP, the Drosophila ortholog of human Cytoplasmic Polyadenylation Element Binding Protein (hCPEB) 2-4 protein subfamily, binds to hundreds of maternally provided, rare-codon-enriched mRNAs in early embryos; that ORB2 targets are translationally repressed and unstable during the MZT; identify a U-rich motif enriched in ORB2 targets’ 3ʹUTRs; and show that this motif confers ORB2 binding and repression to a luciferase reporter mRNA in S2 tissue culture cells. When tethered to a luciferase reporter, ORB2 and hCPEB2 (but not ORB and hCPEB1) repress translation; the C-terminal Zinc-binding (‘ZZ’) domain of ORB2 is necessary and sufficient for repression. ORB2 interacts with a suite of post-transcriptional regulators in early embryos; a subset of these interactions is lost upon deletion of the ZZ domain, notably with the Cup repressive complex. ORB2-targets significantly overlap with those previously identified for the repressive RBP, Smaug (SMG). Analysis of the early embryo’s translatome in the presence or absence of the endogenous ZZ domain shows that mRNAs bound by ORB2 but not by SMG move onto polysomes upon ZZ domain deletion whereas co-bound transcripts do not, consistent with co-regulation of the latter set of transcripts by both RBPs. Our results assign a function to the ZZ domain and position ORB2 in the post-transcriptional network that regulates maternal transcripts during the Drosophila MZT.

**ARTICLE SUMMARY:** We show that Drosophila ORB2, the ortholog of the human CPEB2 RNA-binding protein, negatively regulates its target mRNAs during the maternal-to-zygotic transition via its C-terminal Zinc-binding (‘ZZ’) domain.

## INTRODUCTION

All animal embryos undergo a maternal-to-zygotic transition (MZT) that encompasses two global processes: First, a large subset of maternal gene products loaded into the egg by the female parent is cleared; second, the zygotic genome becomes transcriptionally active (Tadros and Lipshitz 2009; Vastenhouw *et al*. 2019; Harrison *et al*. 2023). In Drosophila embryos, the MZT is both very rapid – reaching completion about 3 hours after fertilization – and extensive: Thousands of maternally loaded transcripts are cleared (De Renzis *et al*. 2007; Lécuyer *et al*. 2007; Tadros *et al*. 2007; Thomsen *et al*. 2010); hundreds of these transcripts are translationally repressed (Kronja *et al*. 2014); and thousands of genes are transcribed from the embryo’s own genome in a process known as zygotic genome activation (ZGA) (Harrison *et al*. 2023). Several RBPs have been shown to function in maternal transcript repression and degradation in *Drosophila*: Smaug (SMG) directs repression and clearance of a subset of maternal transcripts prior to ZGA (Tadros *et al*. 2007; Chen *et al*. 2014); Brain tumor (BRAT) binds to and represses a distinct subset of maternal mRNAs both before and after ZGA (Laver *et al*. 2015); Pumilio (PUM) co-binds a subset of the BRAT targets but primarily directs transcript clearance after rather than before ZGA (Thomsen *et al*. 2010; Laver *et al*. 2015; Haugen *et al*. 2024). In contrast to these ‘repressive’ RBPs, Rasputin (RIN), the Drosophila ortholog of vertebrate G3BP, upregulates the stability and translation of its targets during the MZT (Laver *et al*. 2020a).

Here we focus on the ORB2 RBP, which is of particular interest because it and its vertebrate homologs have been reported to mediate both negative and positive post-transcriptional regulatory functions in a variety of organisms and cell types. ORB and ORB2 are the fly homologs of the vertebrate Cytoplasmic Polyadenylation Element Binding (CPEB) family of RBPs, which canonically interact with U-rich motifs known as cytoplasmic polyadenylation elements (CPEs) in the 3ʹUTR of target transcripts (Hake and Richter 1994). Vertebrates have four CPEBs (CPEB1-4), which can be divided into two groups based on sequence similarity: the CPEB1-like subfamily (containing vertebrate CPEB1 and its homologs, such as Drosophila ORB) and the CPEB2-like subfamily (containing vertebrate CPEB2-4 and their homologs, such as Drosophila ORB2) (Wang and Cooper 2009; Fernández-Miranda and Méndez 2012; Ivshina *et al*. 2014; Duran-Arqué *et al*. 2022).

CPEB proteins are known to function in the regulation of cytoplasmic mRNAs involved in egg maturation, embryonic stem cell division, memory formation, synaptic plasticity, and gametogenesis (Hake and Richter 1994; Groisman *et al*. 2001; Keleman *et al*. 2007; Standart and Minshall 2008; Krüttner *et al*. 2012; Xu *et al*. 2012). Xenopus CPEB1, for example, is a major specificity factor of an mRNA masking complex that represses translation in oocytes and mediates translational activation of its target mRNAs upon egg activation (Hake and Richter 1994; Kim and Richter 2006).

ORB2 has been shown to have a role in neural development, long-term memory formation, and spermatogenesis (Keleman *et al*. 2007; Hafer *et al*. 2011; Majumdar *et al*. 2012; Xu *et al*. 2012; Khan *et al*. 2015; Hervás *et al*. 2016). The *orb2* gene encodes two protein isoforms, ORB2A and ORB2B, which have 542 amino acids in common (from N-to-C): a poly-glutamine (polyQ) tract, a predicted intrinsically disordered region (IDR2) that spans approximately 200 amino acids, the RNA-binding domain (RBD) comprised of tandem RNA recognition motif (RRM) domains, and a ZZ-class Zinc-binding domain. ORB2A and ORB2B differ only in IDR1 at their N-termini (ORB2B is diagrammed in Figure 1a). ORB2A-ORB2B hetero-oligomers serve as translational activators whereas, in the absence of ORB2A, ORB2B does not readily oligomerize and has been shown to repress translation of reporter transcripts *in vitro* and in S2 tissue culture cells (Mastushita-Sakai *et al*. 2010; Khan *et al*. 2015; Stepien *et al*. 2016; Hervas *et al*. 2020). The early Drosophila embryo is unusual in that only one of the two protein isoforms, ORB2B, is expressed, affording us the opportunity to assess ORB2B’s mechanism of action and function *in vivo* in the absence of ORB2A.

**Figure 1.**
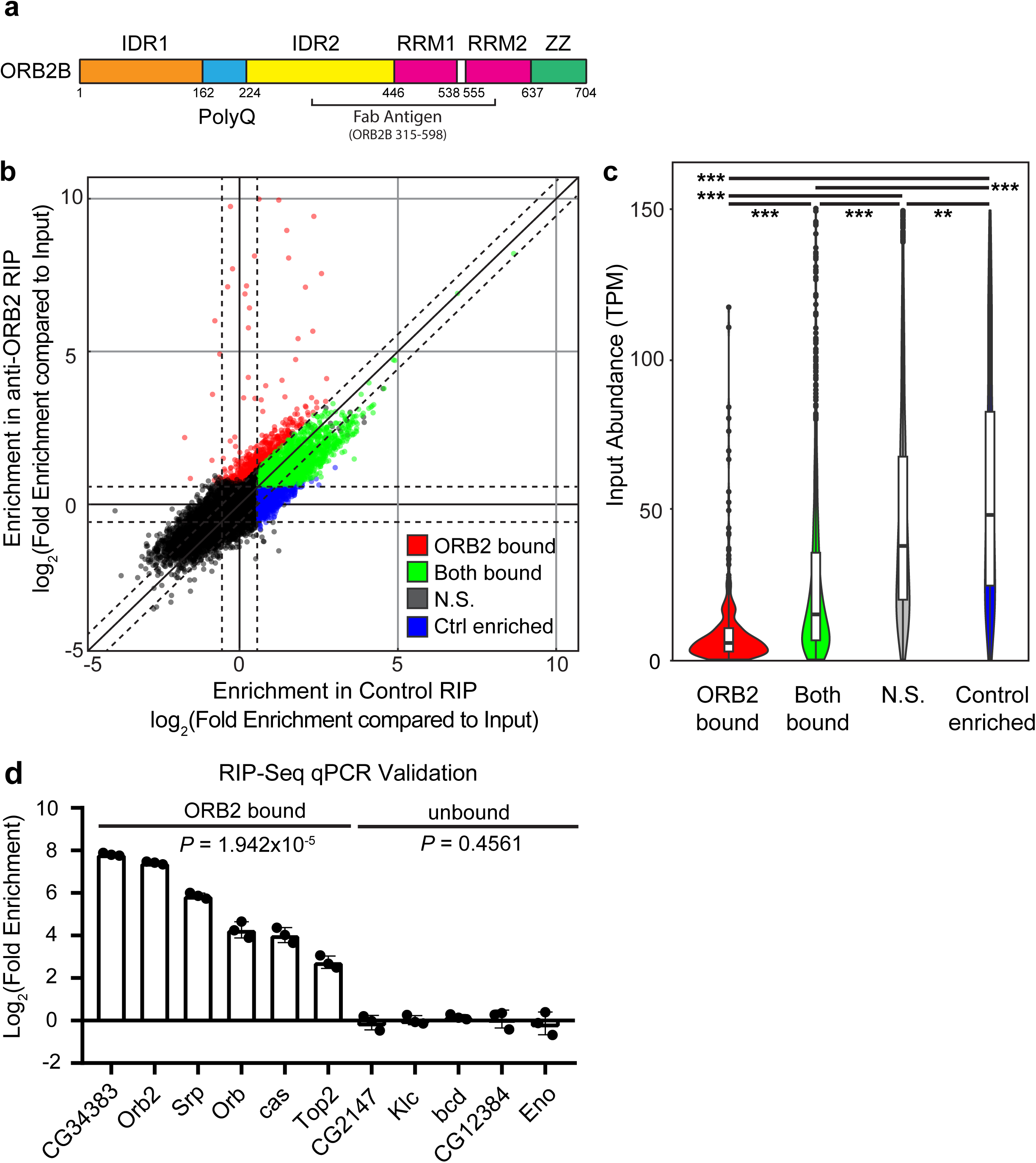
RIP-seq identifies over 450 ORB2 targets in 0-3 hr embryos. (a) Domain structure of ORB2B and location of the antigen used to screen for phage-displayed synthetic antibodies. (b) Scatterplot depicting the Log_2_(Fold Enrichment compared to Input) for all genes identified in the RIP-seq analysis; each point represents a single gene. Values represent the average enrichment from three independent biological replicates. ORB2 targets (red) are defined as having fold enrichment > 1.5 in the anti-ORB2 RIP-seq compared to input and a ratio of ORB2/Control > 1.5, with an FDR adjusted *P*-value < 0.05; genes enriched in both ORB2 and control RIP-seq (green) are defined as having fold enrichment > 1.5 in both anti-ORB2 and control RIP-seq, with a ratio of ORB2/Control < 1.5 and an FDR adjusted *P*-value < 0.05; Control enriched transcripts (blue) are defined as having fold enrichment < -1.5 in the control RIP-seq, fold enrichment < -1.5 in the anti-ORB2 RIP-seq, and an FDR adjusted *P*-value < 0.05. Dotted lines indicate ±1.5 fold-change thresholds. (c) Violin and boxplots depicting the distribution of transcript abundance as measured by average TPM from three biological replicates based on target status as described in (A). Wilcoxon Rank Sum Test with Bonferroni multiple-test correction: ** = *P* < 10^-10^; *** = *P* < 10^-15^. (d) Validation of RIP-seq targets by RT-qPCR; bar graphs depict the log_2_(fold enrichment) in an anti-ORB2 RIP-RT-qPCR compared to control RIP-RT-qPCR; *P-*values given are derived from Wilcoxon Signed Rank Test comparing ORB2 RIP-RT-qPCR to control RIP-RT-qPCR.

To identify protein and RNA components of ORB2B-containing ribonucleoprotein complexes (RNPs) during the MZT, we used a synthetic antibody to immunoprecipitate (IP) endogenously expressed ORB2-containing RNPs followed by either next-generation sequencing of co-immunoprecipitated transcripts (RIP-seq) or mass spectrometry of associated proteins (IP-MS). We show that ORB2 binds hundreds of mRNAs that are translationally repressed and degraded during the MZT. These transcripts are enriched for rare codons in their open reading frames and for a U-rich motif in their 3ʹUTRs. We identify over 200 protein binding partners of ORB2, including components of the 43S translation pre-initiation complex (PIC) and the Cup-ME31B-TRAL-BEL repressive complex (hereafter referred to as the Cup repressive complex). We leverage cell culture assays to verify that ORB2 binding to the U-rich motif results in translational repression of a reporter RNA; to map ORB2’s repressive function to the C-terminal ZZ domain; and to show that the repressive function of the ZZ domain is conserved in its subfamily ortholog, human CPEB2 (hCPEB2), but not in members of the other subfamily, ORB or hCPEB1. We produce a CRISPR deletion of the ZZ domain of endogenous ORB2 and show that deletion of the ZZ domain results in loss of interaction with the PIC and the Cup repressive complex and a shift of ORB2’s targets onto polysomes in early embryos. We show that a subset of ORB2’s targets is co-bound by the SMG RBP. Loss of the ZZ domain results in ‘ORB2 only’ targets (i.e., those not co-bound by SMG) moving onto polysomes; in contrast, ORB2-SMG co-bound targets do not. Together our results provide a function for the ZZ domain and position ORB2 within the RBP regulatory network that controls the post-transcriptional fate of maternal mRNAs. We report elsewhere that the endogenous deletion of the ZZ domain leads to defects in spermatogenesis (Low *et al*. 2025) and brain development (Hailstock *et al*. 2025).

## MATERIALS AND METHODS

### Drosophila stocks

The following *Drosophila melanogaster* stocks were used: *w^1118^* and *orb^ΔZZ^*. The latter stock was generated for use in this study (see below). Flies were cultivated at 25°C under standard laboratory conditions unless otherwise indicated.

### Generation of the CRISPR deletion of the ZZ domain at the ORB2 endogenous locus

Generation of this CRISPR line was performed by Well Genetics (13F.-12, No. 93, Sec. 1, Xintai 5th Rd., Xizhi Dist., New Taipei City 221416, Taiwan (R.O.C.)). The scarless PBac-dsRed method was used to produce the *orb2^ΔZZ^*line (Gratz *et al*. 2013; Gratz *et al*. 2014). The ZZ domain, defined as from D638 to C704 (using the numbering from the ORB2-PB isoform) was targeted for deletion such that the endogenous 3’UTR was retained in its entirety. Two guide RNAs (gRNAs) were used to direct CRISPR to make double-stranded cuts in the DNA surrounding the mutation of interest; in our case, one gRNA was directed to the genome to a cut site just after the beginning of the ZZ domain (-179 nt from STOP Codon of *orb2-RA/B/C/D/H*), and the second gRNA was directed to cut +3 nt from STOP Codon. Cas9 mediates breaks in the double-stranded DNA, which triggers homologous repair between the genomic locus and a donor plasmid, which was co-injected with the gRNA plasmid.

The donor plasmid containing the mutation of interest (a deletion of the ZZ domain, which removes the genomic portion of the coding sequence after the end of the second RRM, advancing the STOP codon to this point, such that this mutant protein would stop at residue L637 in the PB isoform) followed by a dsRed marker flanked by Piggy Bac 5ʹ terminal repeats (Figure S1a). The Piggy Bac transposon terminal repeats enabled this marker to be excised after it had been used to screen for successful CRISPR incorporation into the genome, leaving behind no trace of the marker afterwards (i.e. it is “scarless” on the genome). Flanking this dsRed marker is genomic sequence that is homologous to the base genome, 1kb upstream and downstream for the cut site; this provided the homology needed to facilitate homology-dependent repair (HDR) between the donor plasmid and the genomic site.

After injection of the host *w^1118^* strain (LWG182), balanced fly lines were established from the resulting progeny and screened for successful integration of the mutation-containing region of the donor plasmid. Successful recombination resulted in an ORB2 gene locus harboring a precise deletion of the ZZ domain at the endogenous locus, marked by the expression of dsRed, which was later removed from the genome using the Piggy Bac transposon machinery to excise the marker by crossing them to a Piggy Bac expressing fly (BDSC #8285). PCR was carried out to confirm donor plasmid insertion at the intended location in the correct orientation (Figure S1b) followed by deletion of the dsRed marker (Figure S1c), which was then verified by sequencing (Figure S2). PCR was used to confirm that the *orb2^ΔZZ^* line was not infected with Wolbachia (since BDSC #8285 is known to be infected) (data not shown).

### Synthetic antibody (Fab) expression and harvesting

Phage displayed synthetic antibodies against ORB2 protein have been described previously (Laver *et al*. 2012; Na *et al*. 2016). To summarize, antigens containing amino acids 163-446 (numbered according to ORB2-PA isoform) were expressed and purified from *E. coli* as GST fusion proteins. Synthetic antibody library F (Laver *et al*. 2012; Na *et al*. 2016) was used in five rounds of binding selection, which included negative selection to remove antibodies that bound to GST, to obtain synthetic antibodies that bound to the ORB2 antigen. The two synthetic antibodies used in this study were anti-ORB2 E8 and anti-hEGFR C1 (control).

Synthetic antibodies (‘Fabs’) were expressed and harvested from *E. coli* as previously described (Laver *et al*. 2012). In summary, DH5α *E. coli* was transformed with plasmid containing the FLAG- and 6xHis-tagged Fab open reading frame under the control of the *lac* operator. Transformed cells were used to inoculate bacterial culture using 2xYT media; the inoculated culture was incubated at 37°C on rotation at 225rpm until OD595 reached 0.8. The culture was then induced with isopropyl β-d-1-thiogalactopyranoside (IPTG) to drive expression of the Fab and allowed to grow at 18°C overnight on rotation at 225rpm. Bacteria were pelleted by centrifugation at 4°C, 4000 rpm for 30 minutes, then the bacterial pellet was resuspended with B-PER bacterial protein extraction reagent liquid solution (Thermo Scientific) supplemented with protease inhibitors (150mM NaCl, 1mM AEBSF, 2mM Benzamidine, 2µg/mL Leupeptin, 2µg/mL Pepstatin) and Benzonase nuclease (2500 units/mL). Cells were allowed to lyse for 30 minutes at 4°C, then the lysate was cleared by centrifugation at 4°C at 21,380 RCF for 30 minutes. Supernatant was collected and synthetic antibodies were purified by passing lysates through a column filled with Protein A affinity gel matrix beads; the captured Fab was eluted with an acid buffer containing 50mM NaH_2_PO_4_, 100mM H_3_PO_4_, and 140mM NaCl, pH to 2.8 in eight fractions of 200μL each. Fab-containing fractions were pooled and dialyzed with 1L PBS at 4°C for eight hours, twice. Dialyzed Fab was diluted with an equal volume of 95% glycerol 1xPBS and stored at -20°C.

### Collection of embryo lysates

Lysate was collected from embryos laid and aged at 25°C. Extract was prepared from embryos collected 0 to 3 hours post egg-lay on apple juice agar plates, dechorionated with 100% bleach for 2 minutes, washed with 0.1% Triton X-100, and disrupted by crushing in minimal volume of lysis buffer consisting of 150 mM KCl, 20 mM HEPES-KOH (pH 7.4), 1 mM MgCl_2_, 1 mM DTT, and protease inhibitors (1 mM AEBSF, 2 mM benzamidine, 2 μg/mL pepstatin, 2 μg/mL leupeptin). The lysate was cleared by centrifugation at 4°C at 21,380 RCF for 15 minutes, and the supernatant was collected and either used immediately or stored at -80°C.

### RIP-seq

For each replicate of RIP-seq, 1mL of 0-3 hour embryo extract (prepared as described above) was supplemented with Triton X-100 to 0.1% final concentration, then diluted in an equal volume of lysis buffer with 0.1% Triton X-100 and cleared again by centrifugation at 4°C, 13 000 RPM for 15 minutes. The supernatant was incubated for 3 to 4 hours at 4°C with end-over-end rotation with 100μL of anti-FLAG M2 affinity gel beads (Sigma) which had been pre-incubated at 4°C with 50μg of purified anti-ORB2 E8 Fab blocked with 5μg/μL BSA. After incubation, the lysates were decanted and beads were washed four times with lysis buffer supplemented with 0.1% Triton X-100 and then three times with equilibration buffer: 100 mM NaCl, 100 mM HEPES-NaOH (pH 7.4), 1 mM MgCl_2_. Fabs and associated ribonucleoprotein (RNPs) were eluted from the beads with 200 μL of 200μg/μL FLAG-peptide (Sigma) suspended in equilibration buffer at 4°C for 20 minutes. RNA was isolated from the eluate using TRI Reagent (Sigma) according to the manufacturer’s instructions.

RNA was prepared for sequencing using Ribo-Zero Gold rRNA depletion kit and TruSeq Stranded Total RNA Library Prep Kit (Illumina) according to manufacturer’s instructions. Libraries were sequenced using the HiSeq 2500 System, V4 Chemistry (Illumina), with paired-end reads 125nt in length. The raw data are available on GEO (accession number GSE299661).

### Computational analysis of RIP-seq data

Paired-end reads obtained from the sequencing were mapped to the *Drosophila melanogaster* transcriptome r6.23, FB2018_04 obtained from flybase.org (Ozturk-Colak *et al*. 2024) using the Salmon read mapping program (Patro *et al*. 2017). The reference transcriptome was supplied as a FASTA file. The resulting quantification was then analyzed for enrichment in the ORB2 IP compared to Input and control IP compared to Input using the DESeq2 software package for R (Love *et al*. 2014). Prior to formal analysis, the dataset was filtered for transcripts having > 1TPM in at least three of the six samples, and rlog transformation was performed to stabilize the variance. The similarity between samples was calculated by determining the Poisson Distance between pairs of samples using the *PoiClaClu* package in R (see Figure S3) (Whitten 2011). The differential expression analysis was performed with the *dds* function using alpha = 0.01 cut-off for the False Discovery Rate. This returned a fold enrichment for each transcript, accompanied by an adjusted *P*-value. We defined an ORB2-bound RNA target as satisfying all three of the following criteria: (1) it was enriched in the ORB2 IP >1.5-fold relative to Input; (2) it had an ORB2 IP/Control IP ratio >1.5; (3) it had an adjusted *P*-value < 0.05. The results are presented in Figure 1 and File S1.

For comparisons between our dataset and previously published datasets, each pairwise comparison was done independently. For comparisons that required a Wilcoxon Rank Sum test, for each given pair of datasets, a common background list of genes was first compiled by taking the intersection between the input/background of both datasets; genes which only appeared in one dataset and not the other were discarded. Then, the measurements of the remaining identified genes were placed into a dataframe in which each row corresponded to one unique gene, the first column contained the measurements taken from one dataset, and the second column contained measurements from the other dataset. This dataframe was then used to perform a Wilcoxon Rank Sum test using the wilcox.test(x, y, exact = FALSE) function in R, where x is the data in the first column, y is the data in the second column. The results are shown in Figure 2.

**Figure 2.**
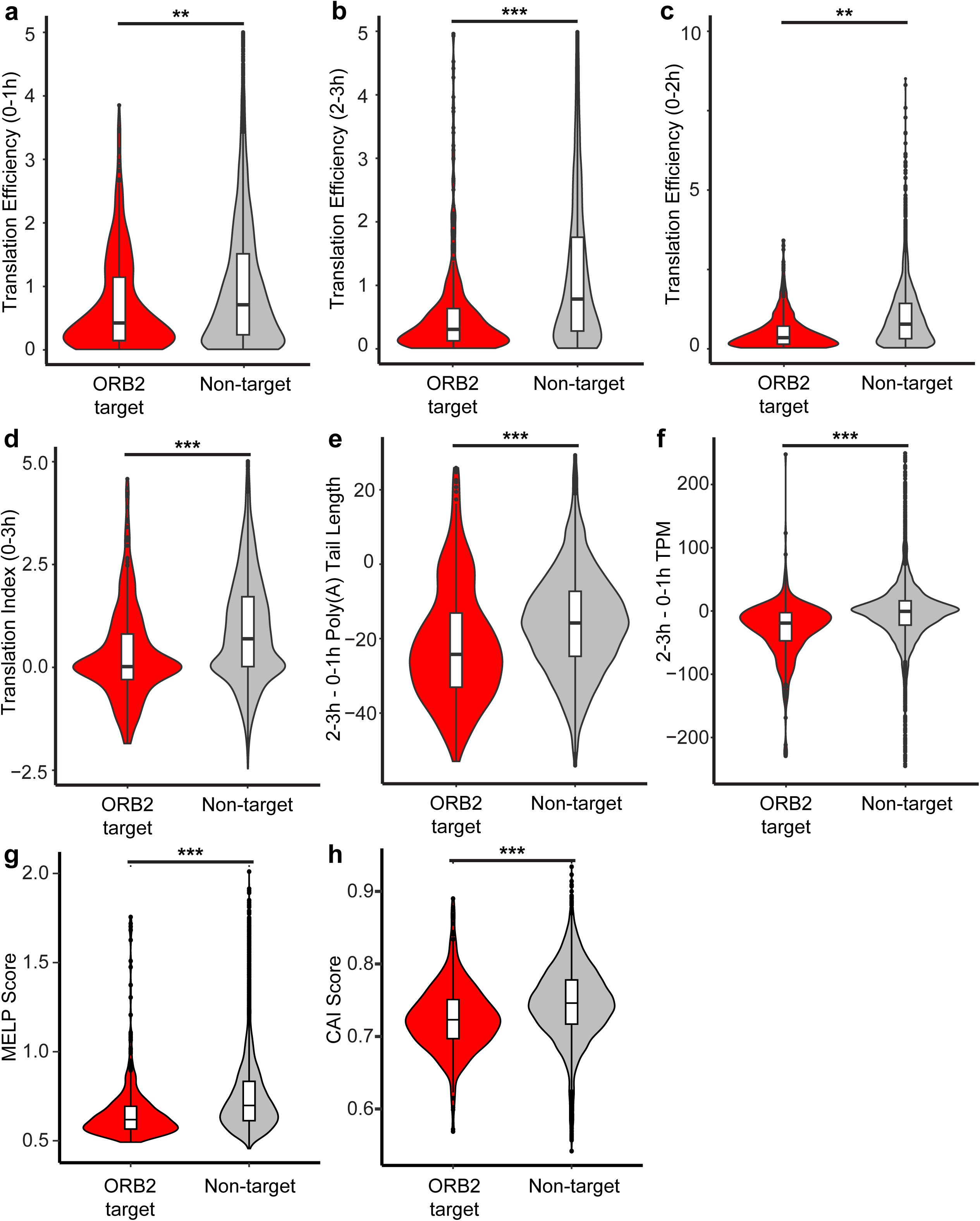
ORB2 target transcripts are enriched in translationally repressed, unstable, rare-codon containing transcripts during the MZT. Violin plots and boxplots depicting the relationship between ORB2-association and translational status, change in poly(A) tail length, and change in transcript abundance. Non-target genes include all expressed genes that were not defined as ORB2 targets in the RIP-seq analysis. (a) Translation efficiency, calculated from ribosome footprinting performed on 0-1 hour embryo extracts (Eichhorn *et al*. 2016). (b) Translation efficiency, calculated from ribosome footprinting performed on 2-3 hour embryo extracts (Eichhorn *et al*. 2016). (c) Translation efficiency as measured by ribosome profiling performed on 0-2 hour embryo extracts (Dunn *et al*. 2013). (d) Translation index, calculated from polysome gradients performed on 0-3 hour embryo extracts (Chen *et al*. 2014). (e) Change in transcript poly(A) tail length, calculated by comparing 0-1 hour and 2-3 hour embryo extracts, as measured by PAL-Seq (Eichhorn *et al*. 2016). (f) Change in abundance, calculated by comparing 0-1 hour and 2-3 hour embryo extracts, as measured by RNA-Seq (Eichhorn *et al*. 2016). (g) MELP score, calculated for all annotated CDS longer than 80 codons on FlyBase 6.58 by coRdon with ribosomal proteins as the references for codon optimization, and the isoform with minimum score (the worst expressivity) is chosen to represent each expressed gene. (h) CAI score, calculated for all annotated CDS longer than 80 codons on FlyBase 6.58 by local version of CAIcal with the *Drosophila melanogaster* codon usage table from the Kazusa codon usage database, and the isoform with minimum score (the most enriched for rare codons) is chosen to represent each expressed gene. Wilcoxon Rank Sum Test: ** = *P* < 10^-10^; *** = *P* < 10^-15^.

For comparisons that required a Fisher’s Exact Test, for each given pair of datasets a common background list of genes was first compiled by taking the intersection between the input/background of both datasets; genes that only appeared in one dataset and not the other were discarded. Then, the remaining identified genes of interest within in each dataset were compared to look for common elements; genes that were identified in both lists of interest were annotated as “Overlap”, whereas genes that only appeared in one list were annotated as “X Only”. A Fisher’s Exact Test was performed by using a 2x2 contingency table, where first cell (top left) contained the number of genes that did not appear in either list (equal to the total number of genes in the common background minus the number of genes described in the other three cells), the second cell (top right) contained number of genes that appeared only on the ORB2 list (ORB2 Only), the third cell (bottom left) contained the number of genes that appeared only on the other list of interest (Other Only), and the last cell (bottom right) contained the number of genes appear on both lists of interest (Overlap). This contingency table was constructed and evaluated using the fisher.test() function in R. The results are presented in Table 1.

**Table 1.**
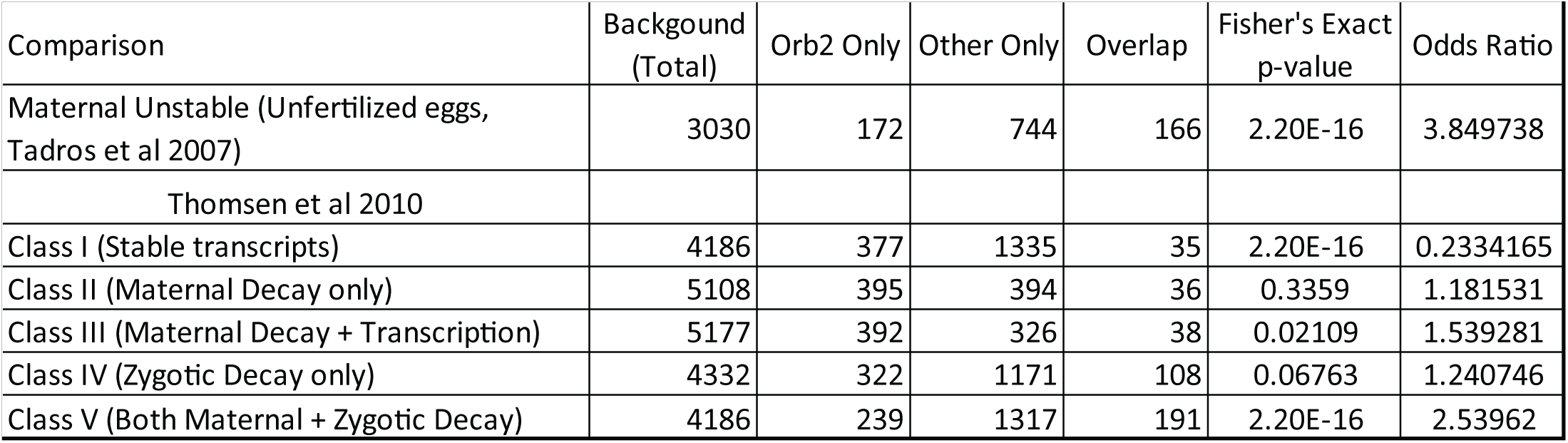
Overlap between the list of ORB2 targets and lists of genes associated with mRNA stability or decay in the early embryo.

The Gene Ontology (GO) term enrichment analysis was performed using the Functional Profiling function of the web version of gProfiler2 (accessed at https://biit.cs.ut.ee/gprofiler/gost) (Kolberg *et al*. 2023). The list of ORB2-bound target genes was provided to the program in the Query field, and the Run Query option was selected in order to begin analysis. The results are presented in Tables 2-4.

**Table 2.** Molecular Function GO-terms enriched in ORB2 targets. GO-term analysis was performed using the gProfiler2 tool (Kolberg *et al*. 2023).

| GO Term Name | Adjusted p-value | GO Term Size | ORB2 targets | Overlap |
| --- | --- | --- | --- | --- |
| protein binding | 1.13E-21 | 3760 | 416 | 229 |
| mRNA 3'-UTR binding | 5.78E-11 | 48 | 416 | 17 |
| mRNA binding | 1.15E-10 | 238 | 416 | 35 |
| mRNA regulatory element binding translation repressor activity | 2.51E-08 | 18 | 416 | 10 |
| translation repressor activity | 7.18E-08 | 25 | 416 | 11 |
| translation regulator activity | 2.28E-04 | 106 | 416 | 16 |
| catalytic activity, acting on DNA | 2.49E-04 | 147 | 416 | 19 |
| protein serine/threonine kinase activity | 5.25E-04 | 184 | 416 | 21 |
| DNA binding | 6.42E-04 | 1127 | 416 | 69 |
| poly-pyrimidine tract binding | 1.28E-03 | 14 | 416 | 6 |
| adenyl ribonucleotide binding | 1.42E-03 | 846 | 416 | 55 |
| catalytic activity, acting on a nucleic acid | 1.62E-03 | 406 | 416 | 33 |
| ATP binding | 2.44E-03 | 840 | 416 | 54 |
| purine ribonucleotide binding | 2.99E-03 | 1021 | 416 | 62 |
| purine nucleotide binding | 3.63E-03 | 1072 | 416 | 64 |
| adenyl nucleotide binding | 3.73E-03 | 896 | 416 | 56 |
| ribonucleotide binding | 4.31E-03 | 1033 | 416 | 62 |

**Table 3.** Biological Process GO-terms enriched in ORB2 targets. GO-term analysis was performed using the gProfiler2 tool (Kolberg *et al*. 2023).

| GO Term Name | adjusted<br>p-value | GO Term<br>Size | ORB2<br>targets | Overlap |
| --- | --- | --- | --- | --- |
| purine ribonucleoside triphosphate binding | 4.48E-03 | 1012 | 416 | 61 |
| anatomical structure development | 5.77E-28 | 2775 | 435 | 212 |
| developmental process | 2.30E-27 | 2954 | 435 | 219 |
| negative regulation of cellular process | 1.73E-25 | 1520 | 435 | 144 |
| cell development | 6.92E-25 | 1450 | 435 | 139 |
| multicellular organism development | 8.26E-25 | 2276 | 435 | 182 |
| cell differentiation | 3.46E-24 | 1768 | 435 | 155 |
| cellular component organization | 1.07E-21 | 3178 | 435 | 217 |
| regulation of metabolic process | 1.81E-21 | 2253 | 435 | 174 |
| gamete generation | 1.19E-20 | 917 | 435 | 100 |
| germ cell development | 7.67E-20 | 721 | 435 | 86 |
| anatomical structure morphogenesis | 1.34E-19 | 1580 | 435 | 136 |
| oogenesis | 1.45E-19 | 582 | 435 | 76 |
| system development | 5.09E-19 | 1527 | 435 | 132 |
| multicellular organismal reproductive process | 1.60E-18 | 1096 | 435 | 107 |
| regulation of primary metabolic process | 1.69E-17 | 1860 | 435 | 146 |
| nervous system development | 4.18E-17 | 1196 | 435 | 110 |
| animal organ development | 5.43E-17 | 1200 | 435 | 110 |
| regulation of macromolecule biosynthetic process | 1.09E-15 | 1742 | 435 | 136 |

**Table 4.**
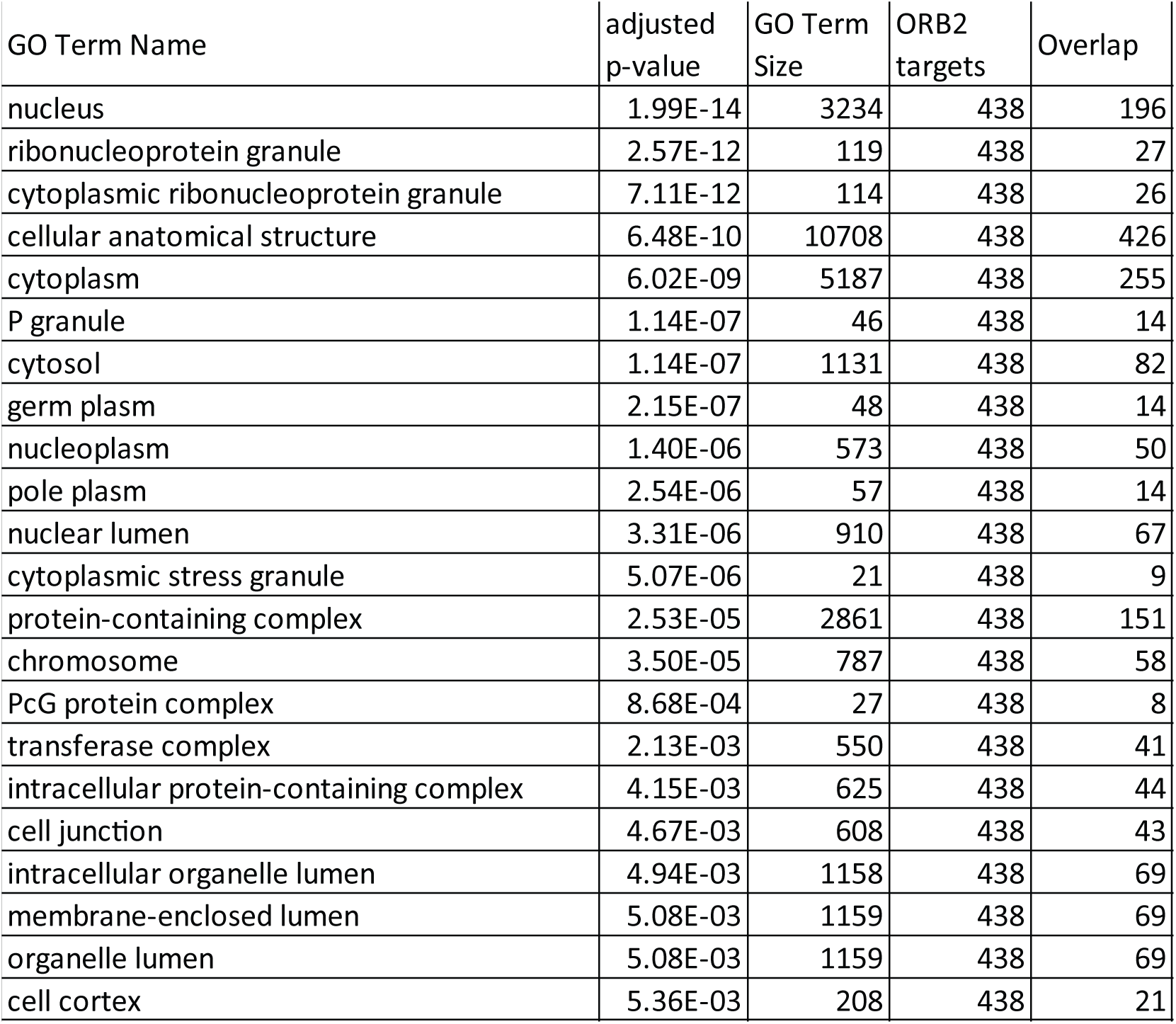
Cellular Component GO-terms enriched in ORB2 targets. GO-term analysis was performed using the gProfiler2 tool (Kolberg *et al*. 2023).

| GO Term Name | adjusted<br>p-value | GO Term<br>Size | ORB2<br>targets | Overlap |
| --- | --- | --- | --- | --- |
| nucleus | 1.99E-14 | 3234 | 438 | 196 |
| ribonucleoprotein granule | 2.57E-12 | 119 | 438 | 27 |
| cytoplasmic ribonucleoprotein granule | 7.11E-12 | 114 | 438 | 26 |
| cellular anatomical structure | 6.48E-10 | 10708 | 438 | 426 |
| cytoplasm | 6.02E-09 | 5187 | 438 | 255 |
| P granule | 1.14E-07 | 46 | 438 | 14 |
| cytosol | 1.14E-07 | 1131 | 438 | 82 |
| germ plasm | 2.15E-07 | 48 | 438 | 14 |
| nucleoplasm | 1.40E-06 | 573 | 438 | 50 |
| pole plasm | 2.54E-06 | 57 | 438 | 14 |
| nuclear lumen | 3.31E-06 | 910 | 438 | 67 |
| cytoplasmic stress granule | 5.07E-06 | 21 | 438 | 9 |
| protein-containing complex | 2.53E-05 | 2861 | 438 | 151 |
| chromosome | 3.50E-05 | 787 | 438 | 58 |
| PcG protein complex | 8.68E-04 | 27 | 438 | 8 |
| transferase complex | 2.13E-03 | 550 | 438 | 41 |
| intracellular protein-containing complex | 4.15E-03 | 625 | 438 | 44 |
| cell junction | 4.67E-03 | 608 | 438 | 43 |
| intracellular organelle lumen | 4.94E-03 | 1158 | 438 | 69 |
| membrane-enclosed lumen | 5.08E-03 | 1159 | 438 | 69 |
| organelle lumen | 5.08E-03 | 1159 | 438 | 69 |
| cell cortex | 5.36E-03 | 208 | 438 | 21 |

### Rare-codon enrichment analysis

#### MELP score (Supek and Vlahovicek 2005)

Assuming ribosomal proteins are the best translated and the most codon optimized genes in a given genome, a chi-square based statistic called Measure Independent of Length and Composition (MILC) was calculated to evaluate the difference of codon usage frequencies between ribosomal proteins and all annotated CDS that are longer than 80 codons from Flybase 6.58. Then, MELP (MILC-based Expression Level Predictor) was calculated based on MILC to predict expression level for each mRNA, and the isoform with the minimum score (the worst predicted expression) was chosen to represent each expressed gene (File S2). The Wilcoxon Rank Sum Test was performed for protein coding genes that were either ORB2 target or non-target.

#### CAI (Puigbo *et al*. 2008)

Codon Adaptation Index (CAI) is another metric that evaluates the similarity of codon usage to a reference set. This score was calculated for all annotated CDS longer than 80 codons from Flybase 6.58 with the local version of CAIcal and the *Drosophila melanogaster* codon usage table from the Kazusa codon usage database (Nakamura *et al*. 2000). The isoform with the minimum score (the most enriched for rare codons) was chosen to represent each expressed gene (File S2), and the Wilcoxon Rank Sum Test was performed for protein coding genes that were either ORB2 target or non-target.

### Number of Accessible Target Sites (#ATS) *de novo* motif discovery

The ORB2 motifs were identified using the #ATS model, as described (Li *et al*. 2010). Briefly, the #ATS program is a *de novo* motif discovery algorithm which takes as input nucleic acid sequences belonging to mRNA transcripts in two lists: a positive (in this case, the 3ʹUTR of ORB2 targets) and negative (in this case, the 3ʹUTR of co-expressed and unbound) list, and outputs *k*-mers that best differentiate the positive transcripts from the negative transcripts. To begin the analysis, 6-nt long seed sequences are first assessed based on their enrichment in the positive list compared to the negative list, and then the top seed sequences are used as the lauching point upon which the program iterates in an attempt to improve enrichment scores.This is assessed based on enrichment in the positive list compared to the negative list and accounts for RNA accessibility, as predicted by computational modeling of RNA secondary structures, and assigned a *P*-value (see File S3). Iterative enrichment yielded degenerate motifs, so the top five seed 6-mers from the #ATS were combined into a consensus sequence by aligning the G in the motif and converting the resulting table into a position weight matrix (PWM) shown in Figure 3.

**Figure 3.**
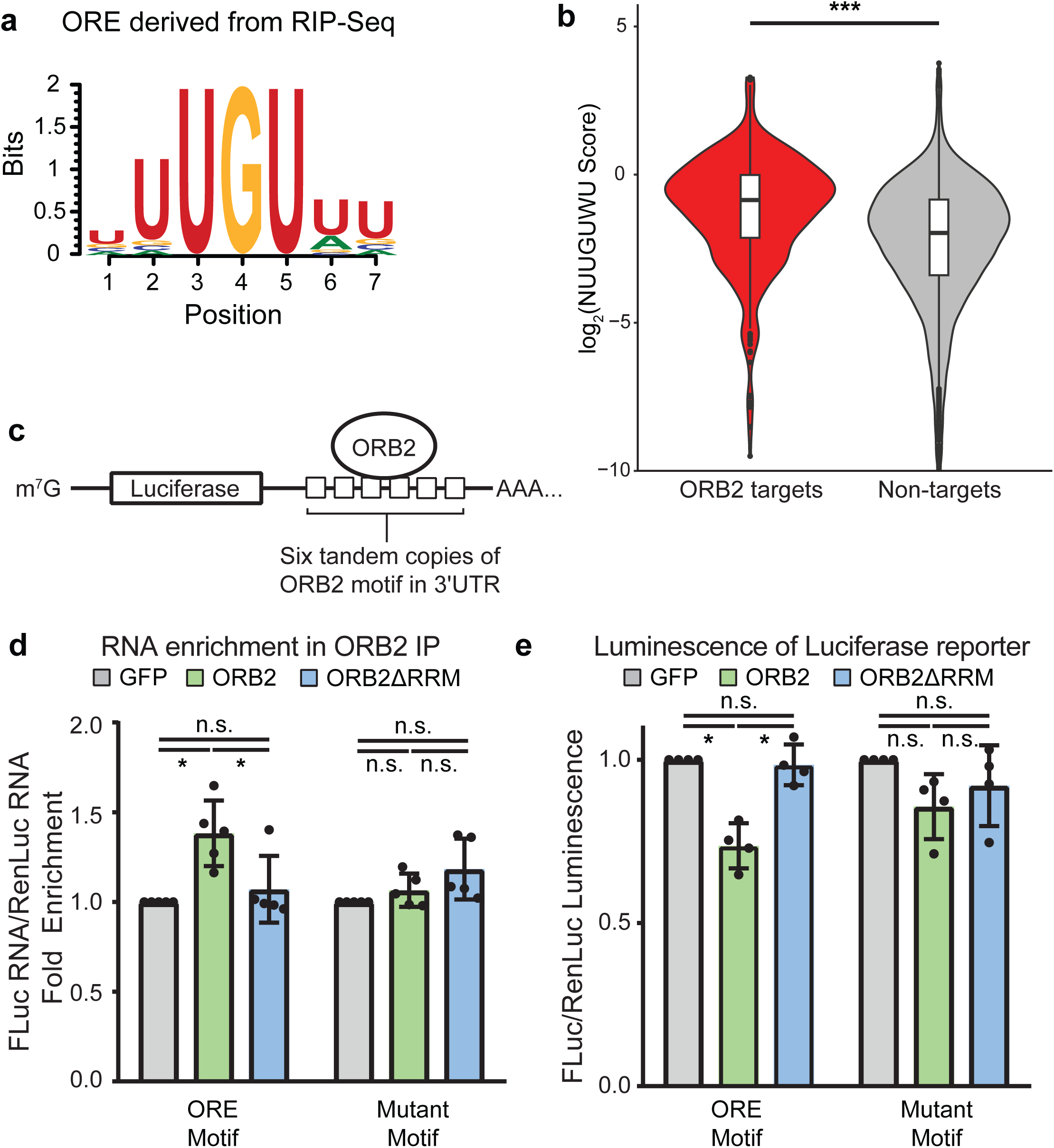
ORB2 binding motifs predicted from RIP-seq confer translational repression on a reporter mRNA in S2 cells. (a) The #ATS *de novo* motif discovery program (Li *et al*. 2010) was performed on the results from the anti-ORB2 RIP-seq in this study. #ATS identified NUUGUWU as the ORB2 recognition element (ORE) enriched in ORB2 targets. (b) The transcriptome was evaluated for accessible enrichment of the NUUGUWU motif and each transcript was scored based on the number and accessibility of these motif sites. ORB2 targets had significantly higher scores compared to non-targets, as assessed by Wilcoxon rank sum test. *** = *P* < 10^-15^ (c) Diagram depicting the reporter firefly luciferase mRNA with six copies of different ORB2-binding motifs in the 3ʹUTR. (d) Bar graph depicting the enrichment of the luciferase reporter mRNA in an anti-ORB2 RIP-qPCR compared to Renilla luciferase mRNA, normalized to a reporter without ORE motifs. Mutation of the OREs or deletion of the RNA-binding domain (RRM) abrogate binding to the mRNA. (e) Bar graph depicting the ratio of luminescence between firefly luciferase and Renilla luciferase. Mutation of the OREs or deletion of the RNA-binding domain (RRM) abrogate repression. One-way Kruskal-Wallis non-parametric test with Dunn multiple test correction: * = *P* < 0.05.

### Evaluation of ORB2 binding motif enrichment in RNA sequences

The enrichment of ORB2 binding motifs was computationally evaluated using a multi-step pipeline modified from our previous method, which assessed Smaug recognition element (SRE) enrichment (Chen *et al*. 2014). Briefly, the computation program scans each RNA using RNAplfold (ViennaRNA package version 2.3.1) using parameters W=170 -L=120 -T=25 (Bernhart *et al*. 2006; Lorenz *et al*. 2011). Next, the nucleotide sequence is scanned for the motif of interest (either the ORB2 motif uncovered by our analysis, NUUGUWU, or the ORB2 motif determined by RNA Compete, KUUUKKK) (Ray *et al*. 2013). At candidate motif sites, the program then assesses the accessibility of the RNA at that position using the results from the RNAplfold analysis. A score is then assigned to each motif site based on the probability that the motif sequence is unpaired with any other nucleotide in the sequence. The total score for a transcript is represented by the sum of motif scores for all motifs found in the sequence.

### Cell culture and transient transfection

The Drosophila S2 cell line was maintained at 25°C with ExpressFive SFM (Fisher Scientific) medium supplemented with 16mM L-glutamine, 100units/mL penicillin, and 100µg/mL streptomycin. 400µL of S2 cells at a density of 1.75×10^6^ were plated into one well of a 24-well tissue culture tray and transfected with a plasmid mixture containing 1.5ng firefly luciferase plasmid, 1.5ng Renilla luciferase plasmid, 20ng FLAG-effector plasmid (containing the open reading frame for either ORB2 whole protein, fragments of ORB2 protein, Protein A, or GFP), 4ng of FLAG-GST plasmid, and 178.5ng pSP72 plasmid using 0.4µL of TransIT-Insect transection reagent (Mirius Bio) suspended in 20µl of tissue culture medium according to the manufacturer’s instructions. For tethering assays, the luciferase plasmid was modified such that the luciferase 3ʹUTR contained six copies of the TAR element and the effector protein construct was modified to include the BIV-Tat ORF N-terminal to the FLAG tag. For experiments which measure ORB2 binding to ORE elements, the luciferase plasmid was modified such that the luciferase 3ʹUTR contained six copies of either intact ORE elements (UUUUGUU) or mutant ORE elements (UUAUCUC). Both luciferase reporters were under the control of the metal-inducible metallothionein promoter; all FLAG-tagged effector constructs were derived from pAc5.1/V5-His plasmid (Thermo Fisher Scientific), which carries the Actin5C promoter. Luciferase expression was induced 24 hours post-transfection by the addition of copper sulphate to final concentration of 0.5mM. 24 hours post-induction and at 48 hours post-transfection cells were harvested for downstream assays.

### Luciferase Assay

Cell lysate was prepared from S2 cell transient transfections by dislodging cells from the surfaces of the cell culture container and transferred into conical tubes. Cells were pelleted by centrifugation at 21,380 RCF at 4°C for 1 minute; the supernatant was removed, and the cell pellet was resuspended in by adding 20 µL of lysis buffer (containing 1xPassive Lysis Buffer [Promega], 10 mg/mL BSA, and 1 mM AEBSF). Lysis was facilitated by pipetting up and down and allowing the cells to lyse at 4°C for 10 minutes. Firefly and Renilla luciferase activities were measured using the Dual-Luciferase Reporter Assay system (Promega) according to manufacturer’s instructions using a Berthold Detection Systems SIRIUS Luminometer. Normalized luminescence values were calculated by first dividing the Firefly luminescence by the Renilla luminesence readings to obtain a Firefly/Renilla ratio; ratios for each biological replicate were obtained from the average of two technical replicates. Then, this ratio was normalized to the corresponding GFP control sample (transfected in parallel to the experimental conditions). Normalized expression was presented as the average of at least three biological replicates with error bars representing standard deviation.

### Reverse Transcription – Quantitative Real-time Polymerase Chain Reaction (RT-qPCR)

RNA isolated from ORB2 co-immunoprecipitations was reverse transcribed into single-stranded cDNA with the Superscript IV reverse transcriptase kit (Invitrogen) according to manufacturer’s instructions. 500ng of total RNA per sample were used to synthesize cDNA, which was primed using random hexamer primers. The resulting single-stranded cDNA was diluted 1:25 using RNase-free water and used to perform quantitative real-time PCR (qPCR) with primers specific to the various transcripts assayed. Primers were designed using NCBI Primer-BLAST to cover all isoforms produced by a gene and were required to span an exon-exon junction. qPCR was performed with SensiFAST SYBR PCR mix (Bioline) following the manufacturer’s protocol and using 5 µL of diluted cDNA per reaction. The CFX384 Real-Time System (Bio-Rad) was used to carry out the PCR reaction.

The results of the qPCR were analyzed using CFX Manager software (Bio-Rad). Each biological replicate represented a measurement obtained for a separate embryo collection; values from three technical replicates were averaged and relative gene expression was normalized to the ribosomal protein-coding transcript *RpL32*. Normalized expression was presented as the average of three biological replicates with error bars representing standard deviation.

### Western Blots

The following primary antibodies were used: mouse anti-ORB2 4GS (1:50) obtained from the Developmental Studies Hybridoma Bank (https://dshb.biology.uiowa.edu/orb2-4G8) and described in (Hafer *et al*. 2011), mouse M2 anti-FLAG (Sigma, 1:5,000), and mouse anti-Actin (Sigma Cat#A4700, 1:10,000) which was used as a loading control.

Dechorionated embryos were counted, then lysed in Laemmli buffer at a concentration of 1 embryo/μL and boiled for 2 minutes. Proteins were resolved by loading 5μL of lysed embryos in Laemmli buffer per lane into an SDS-PAGE and transferred to PVDF membrane in the absence of SDS. The blot was blocked at room temperature with 2% non-fat milk in PBST for 30 minutes and incubated with primary antibodies diluted in 2% non-fat milk in PBST at 4°C overnight. After this incubation, the blot was washed 3 x 10 minutes with PBST at room temperature and incubated with the appropriate HRP-conjugated secondary antibodies (1:5000, Jackson ImmunoResearch) in 2% non-fat milk in PBST at room temperature for 1 hour. The blot was washed 3 x 10 minutes with PBST and developed using Immobilon Luminata Crescendo Western HRP substrate (Millipore), imaged using a BioRad Chemidoc MP Universal Hood III imager and band intensities were quantified using ImageLab.

### Structure Prediction with AlphaFold

For structural predictions the full amino acid sequence of the ZZ domain constructs used in the tethering assays (see above) was predicted using AlphaFold version 3 (Jumper *et al*. 2021; Abramson *et al*. 2024). Predictions were visualized and truncated to show regions A-to-D in ChimeraX version 1.9 (Meng *et al*. 2023). Structures were color-coded to depict their predicted local distance difference test (pLDDT) score using the AlphaFold Error Plot tool. The results are presented in Figure S8.

### Mass spectrometry

#### Immunoprecipitation of ORB2 from embryo lysates with synthetic antibodies

For IP-MS, 500 µL of 0-3 hour embryo extract (prepared as described above) was supplemented with Triton X-100 to 0.1% final concentration, then diluted in an equal volume of lysis buffer with 0.1% Triton X-100 and cleared again by centrifugation at 4°C at 21,380 RCF for 15 minutes. The supernatant was incubated for 3 to 4 hours at 4°C with end-over-end rotation with 100μL of anti-FLAG M2 affinity gel beads (Sigma) which had been pre-incubated at 4°C with 50μg of purified anti-ORB2 E8 Fab blocked with 5μg/μL BSA. After incubation, the lysates were decanted and beads were washed twice with lysis buffer supplemented with 0.1% Triton X-100, then another twice with lysis buffer without Triton X-100; beads were transferred into a new tube and washed another twice with lysis buffer without Triton X-100. Fabs and associated proteins were eluted by tryptic digest: Beads were resuspended in 200 µL of 50mM ammonium bicarbonate pH 8 supplemented with 2 μg of MS-grade trypsin and incubated at room temperature overnight with end-over-end rotation. The digested peptides were collected and a second elution with 200 μL of 50mM ammonium bicarbonate was performed at room temperature for 30 minutes. The two fractions were pooled, flash frozen with dry ice, dehydrated in a rotary evaporator for 8 hours, then stored at -20°C.

### Liquid chromatography and mass spectrometry

For liquid chromatography (LC) the following solvents were used: Solvent A (0.1% (vol/vol) formic acid) and Solvent B (0.1% (vol/vol) formic acid/90% (vol/vol) acetonitrile). The LC parameters were set using an Easy-nLC 1200 instrument. Peptides were loaded and separated on a nanoViper trap column (75μm x 2 cm), Acclaim PepMap 100 (C18, 3μm, 100 Å, Thermo Scientific, cat. no. 164946), and EASY-Spray analytical column (75μm x 50cm) Acclaim PepMap RSLC (C18, 2μm, 100Å, Thermo Scientific) using a flow rate of 225nL/minute. Tandem MS was performed using the Q Exactive HF-Orbitrap mass spectrometer (Thermo Scientific), as previously described (Liu *et al*. 2014; Jiang *et al*. 2015; Chiu *et al*. 2016). The parameters for acquisition were 1 MS scan (50 ms; mass range, 390 to 1800) at a resolution of 60,000K, followed by up to 20 MS/MS scans (50 ms each) at a resolution of 15,000 and an AGC target of 1x105. Candidate ions with charge states +2 to +7 were isolated using a window of 1.4 amu with a 5s dynamic exclusion window.

### Data processing and analysis

Four biological replicates were performed, both in the presence or absence of RNase A. The ProHits software package was used to perform Significance Analysis of INTeractome (SAINT) in order to identify ORB2-interacting proteins (Liu *et al*. 2010; Choi *et al*. 2012; Liu *et al*. 2012). Proteins with associated peptide counts were filtered through the Trans Proteomic Pipeline (TPP) for iProphet probability > 0.95 and number of unique peptides ≥ 2. SAINT express analysis was performed on the ProHits interface with only Drosophila detected proteins (selected options: number of compressed controls = 3, burn-in period, nburn = 2000; iterations, niter = 5000, lowMode = 1; minFold = 1; normalize = 1; nCompressBaits = 3). This analysis determined the probability of interaction for each identified prey protein. Proteins with AvgP score (‘SAINT score’) ≥ 0.95 and BFDR ≤ 0.01 were defined as ORB2-ineracting proteins. Raw data are available at MassIVE MSV000098275. The results are presented in File S4.

To perform network analysis on the list of identified protein interactors, we used the GeneMANIA application (v3.5.2) (Warde-Farley *et al*. 2010; Franz *et al*. 2018) with the Cytoscape network visualization program (v3.9.1) (Shannon *et al*. 2003; Montojo *et al*. 2010). Data resulting from the IP-MS was imported to Cytoscape in table form. GeneMANIA maintains a database of functional association data, which includes protein interactions, genetic interactions, pathways, co-expression, and co-localization data that is obtained from a wide variety of published sources. The details regarding the GeneMANIA database can be found online (https://genemania.org/data/). Two interaction networks were produced by performing a Network Search on a subset of identified protein interactors using GeneMANIA’s *Drosophila melanogaster* functional association database (see https://genemania.org/data/current/Drosophila_melanogaster/ for details). Then, the nodes of each network were annotated according to the association type reported in the GeneMANIA database (i.e. known physical interaction, known genetic interaction, or predicted interaction). The results are presented in Figure 5c,d.

**Figure 4.**
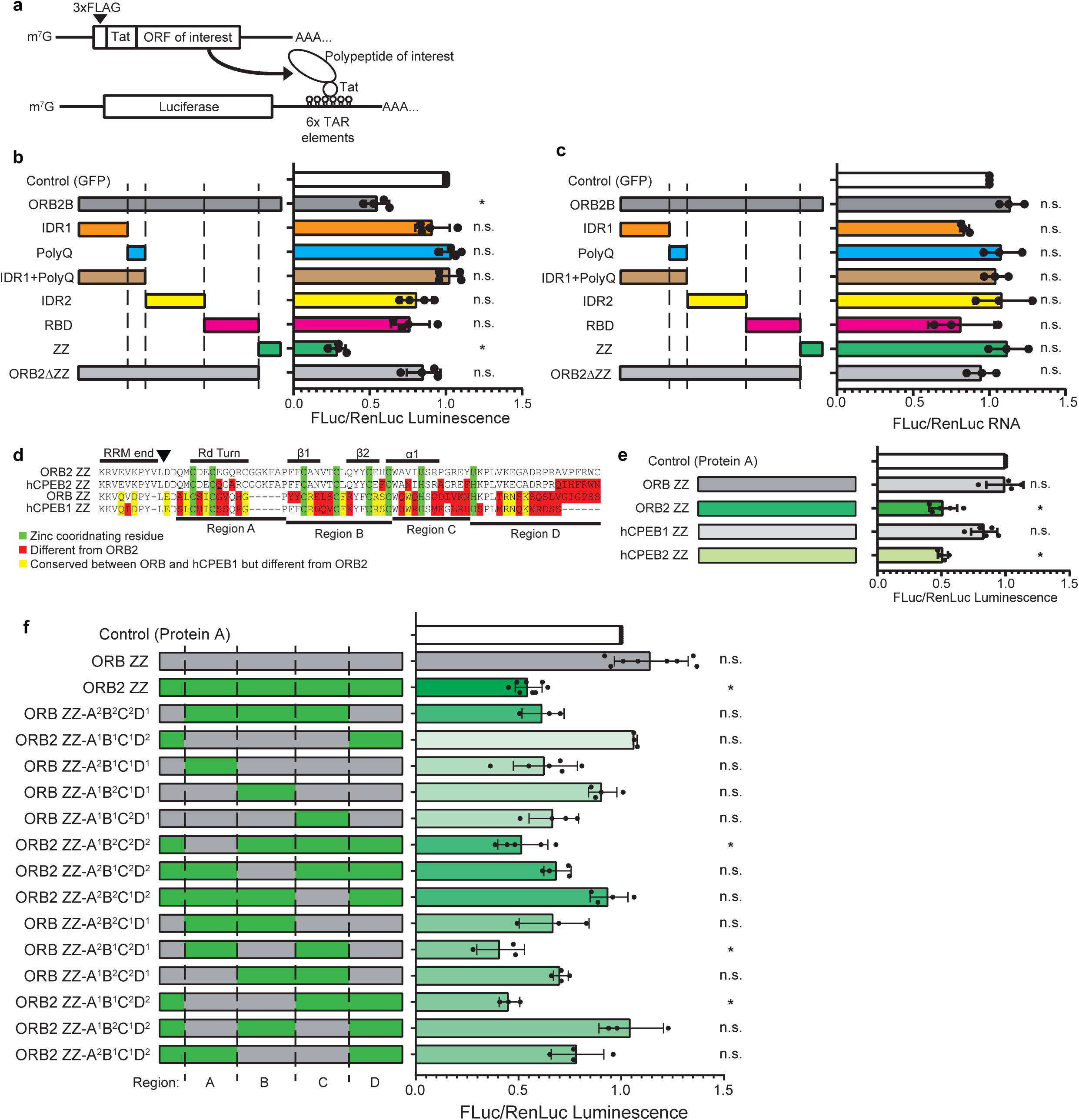
The ORB2 and hCPEB2 ZZ domain but not the ORB and hCPEB1 ZZ domain repress a luciferase reporter. (a) Diagram depicting the tethering assay in which a polypeptide of interest fused to the BIV-Tat domain is tethered to a firefly luciferase reporter mRNA carrying six tandem repeat TAR elements in the 3ʹUTR. (b) Bar graph depicting the expression of the reporter firefly luciferase (normalized to co-transfected Renilla luciferase and normalized to steady state RNA levels) when tethering whole ORB2 or ORB2 fragments compared to GFP control. (c) Bar graph depicting the expression of the reporter firefly luciferase mRNA measured by RT-qPCR (normalized to co-transfected Renilla luciferase mRNA) when tethering whole ORB2 or ORB2 fragments compared to GFP control. Reporter mRNA levels are unaffected by ORB2 ZZ or ORB2ΔZZ. (d) amino acid sequence alignment of the ZZ domains of ORB2, ORB, and their human orthologs, hCPEB1 and hCPEB2. The arrowhead denotes the junction between the RBD and ZZ domains. (e) Bar graph depicting the expression of the reporter firefly luciferase (normalized to co-transfected Renilla luciferase) when tethering different CPEB ZZ domains. Only ORB2 and hCPEB2 repress. (f) Bar graph depicting the expression of the reporter firefly luciferase when ORB ZZ chimeric domains harboring substitutions from the ORB2 ZZ domain, are tethered. One-way Kruskal-Wallis non-parametric test with Dunn multiple test correction: *= *P* < 0.05.

**Figure 5.**
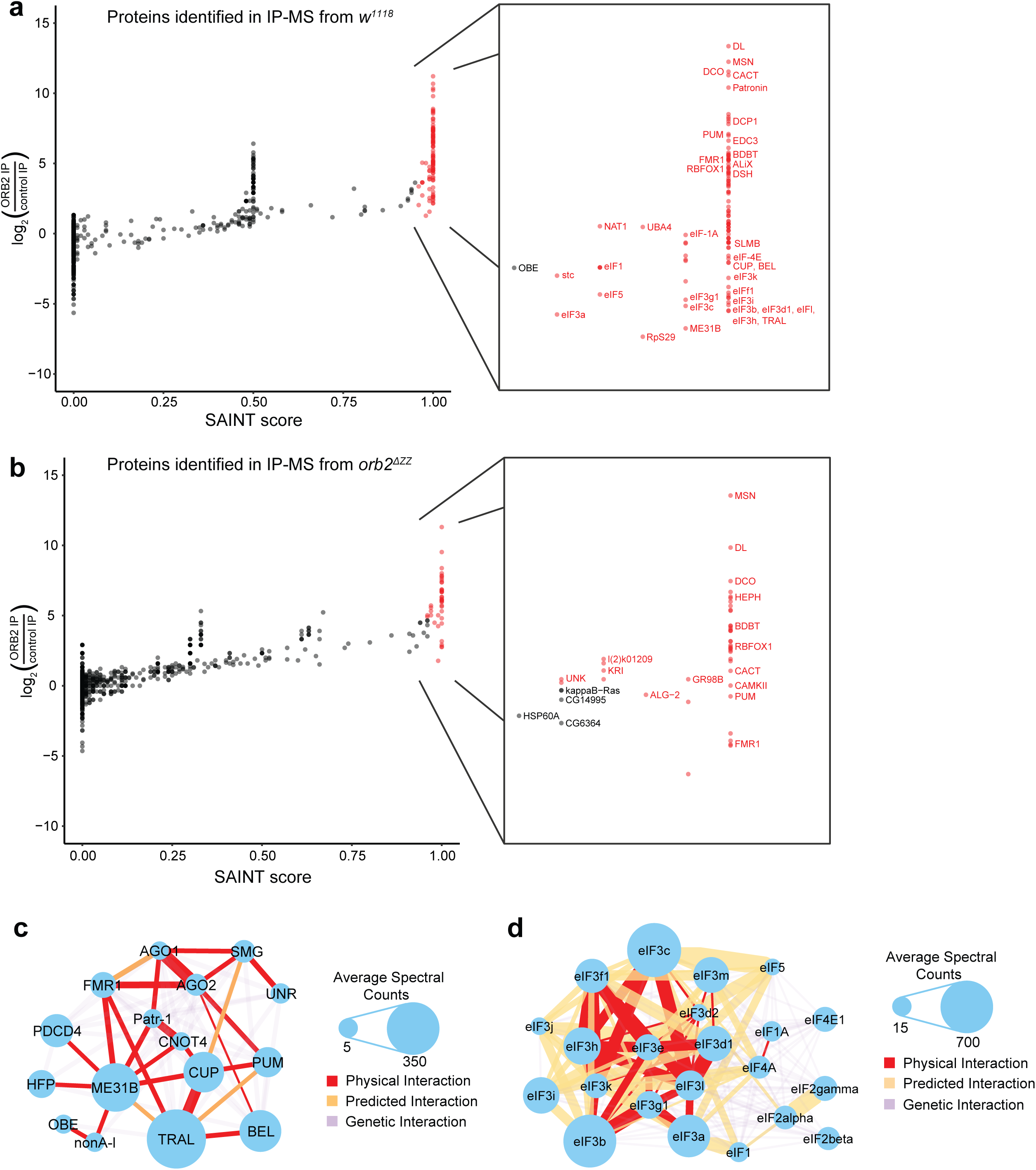
Anti-ORB2 IP-MS from early embryo extracts identifies known translation repressors and the PIC as binding partners. (a-b) Scatterplots depicting all proteins identified in the IP-MS performed in the presence of RNase. Enrichment is expressed as the Log_2_ ratio of the average spectral counts detected in the ORB2 IP compared to the average spectral counts detected in the control IP. Protein interactors were identified by SAINT (with a SAINT score ≥ 0.95 and BFDR < 0.01), shown in red. (a) IP-MS performed on embryos laid by *w^1118^* females, (b) IP-MS performed on embryos laid by *orb2^ΔZZ^* females. (c) Interaction network depicting the connections between components of the Cup repressive complex and other translational repressors identified as ORB2 interactors by SAINT analysis. (d) Interaction network depicting the connections between components of eukaryotic PIC identified as ORB2 interactors by SAINT analysis. Interaction networks were produced by the GeneMANIA app for Cytoscape (Franz *et al*. 2018); size of node indicates average number of spectral counts obtained in the ORB2 IP-MS; thickness of edges indicates strength of interaction assigned by GeneMANIA.

### Embryo sample collection for *orb*^ΔZZ^ time-course and polysome gradient RNA-seq experiments

Embryos were collected from *orb2^ΔZZ^* and *w^1118^* (pre-injected control, LWG182) strains at 1-hour intervals post-egglay (0-1 hour, 1-2 hour, 2-3 hour), dechorionated with ice cold 100% bleach for 2 minutes, washed with 0.1% Triton X-100, and disrupted by crushing using 0.5 g embryos per mL of either lysis buffer (150 mM KCl, 20 mM HEPES-KOH (pH 7.4), 1 mM MgCl2, 1 mM DTT, 1 mM AEBSF, 2 mM benzamidine, 2 μg/mL pepstatin, 2 μg/mL leupeptin) or polysome lysis buffer (7.5 mM MgCl2, 500 mM NaCl, 25 mM Tris (pH 7.5), 1 mM DTT, 1 mM AEBSF, 2 μg/mL leupeptin, 2 mM benzamidine, 2 μg/mL pepstatin A, 2 mg/mL heparin, 0.5 mg/mL cycloheximide). The lysate with lysis buffer was cleared by centrifugation at 4°C at 21,380 RCF for 15 minutes, TRIzol (Invitrogen) was added to the supernatant which used immediately for RNA extraction or stored at -80°C. The lysate with polysome lysis buffer was either used immediately for polysome gradient processing or stored at -80°C.

### Polysome gradients

The polysome gradient protocol was adapted from our published method with modifications (Chen *et al*. 2014). Lysed samples were diluted 1 in 10 in polysome lysis buffer and 20% Triton X-100 was added to a final concentration of 1%, centrifuged at 6,000xg for 10 minutes and the resulting supernatant was diluted in polysome lysis buffer supplemented with 1% Triton X-100 to an A_260_ of 12.5. A 5 ml 15% to 45% linear sucrose gradient in 7.5 mM MgCl_2_, 500 mM NaCl, 50 mM Tris pH 7.5 was produced using a BioComp Model 117 Gradient Mate gradient maker (Biocomp, Fredericton, New Brunswick, Canada) using a rotation angle of 86 degrees and a rotation speed of 25 for 51 seconds. After chilling the polysome gradient at 4°C for 1 hour, 200 μL of diluted embryo extract was loaded onto the top of the gradient, which was then spun at 36,000 rpm in a Beckman SW50.1 rotor for 2.5 hours. 400 μL fractions (hand fractionated) were collected, and A_260_ absorbance was measured using a Nanodrop (Thermo Fisher Scientific). We added 20% SDS, 0.5 M EDTA and 20 mg/ml proteinase K to final concentrations of 0.8%, 0.01 M and 0.128 mg/ml, respectively, and then incubated the fractions for 30 minutes at 37°C. 2.5 µL GlycoBlue (15 mg/mL, ThermoFisher) following which 2.5x volume ethanol was added to each fraction and precipitated overnight. The resulting pellet was washed with 75% ethanol and resuspended in phenol-saturated water.

The fractions were then separated into three pools: Pool 1 contained the (top) fractions 1-3, pool 2 contained fractions 4-6, pool 3 contained fractions 7-13. Following two phenol-chloroform extractions, the aqueous phase of each pool was collected with Invitrogen Phasemaker Tubes (Fisher A33248) following the manufacturer’s protocol. The samples were precipitated by the addition of 7.5 M LiCl to a final concentration of 3.75 M followed by an overnight incubation at -20°C. The pellets were washed with 70% ethanol and resuspended in water.

RNA purification was performed using 2.2× volume NEBNext RNA Sample Purification Beads (NEBNext Ultra II DNA Library Prep with Sample Purification Beads, E7767L) following the manufacturer’s protocol. RNA concentration was assessed with the Qubit RNA HS Assay Kit (Thermo Fisher) and Agilent bioanalyzer at The Centre for Applied Genomics (TCAG), The Hospital for Sick Children, Toronto, ON, Canada.

### rRNA depletion, library preparation and RNA-seq for time-course and polysome gradient RNA-seq

For the RNA-seq time-course rRNA depletion was performed as previously described (Haugen *et al*. 2024). For polysome gradient RNA-seq, rRNA depletion was performed using the QIAseq FastSelect rRNA Fly Kit (QIAGEN, Cat# 33326). Library preparation for RNA-seq was conducted using the NEBNext Ultra II Directional RNA Library Prep Kit for Illumina, and sequencing was performed on an Illumina NovaSeq 6000 (PE150) at the Next Generation Sequencing Facility, The Centre for Applied Genomics (TCAG), The Hospital for Sick Children, Toronto, ON, Canada.

### Computational analysis of time-course and polysome gradient RNA-seq data RNA-seq data processing pipeline

Adapter trimming and quality control used Trim Galore (Version 0.6.5) [https://github.com/FelixKrueger/TrimGalore]. rRNA and ‘unwanted’ RNA removal used SortMeRNA (version 2.1) (Kopylova *et al*. 2012), filtering against *Drosophila melanogaster* 7SLRNA, rRNA, pseudogene-rRNA, tRNA, mtRNA and other species’ rRNA databases. Transcriptome alignment was carried out with Salmon (Version 1.9.1) (Patro *et al*. 2017) in mapping-based mode using FlyBase release 6.23 transcriptome RNA (FB2018_04) (Ozturk-Colak *et al*. 2024). Full decoy indexing was applied following the Salmon website instructions [https://salmon.readthedocs.io/en/latest/salmon.html].

### Differential gene expression analysis

Differential expression analysis used the R package tximeta (version 1.20.3) (Love *et al*. 2020) and Deseq2 (version 1.42.1) (Love *et al*. 2014) in R (version 4.3.2). For the *orb2^ΔZZ^* and wild type time-course total RNA-seq, reads from each time point were normalized using the DESeq2 package. For the *orb2^ΔZZ^* and wild type polysome gradient RNA-seq time-courses, reads of each time point of *orb2^ΔZZ^* and wild type pool 1 and pool 3 RNA were normalized using the DESeq2 package. For each gene, the mean of normalized counts per pool was computed at each time point and genotype. The ‘Translation Index’ of each mRNA was calculated as the ratio = mean normalized counts of pool 3 /mean normalized counts of pool 1. Normalized counts were used for statistical analysis by the Wilcoxon Rank Sum Test and data were visualized with ggplot2 in R. Raw data are available at GEO for the RNA-seq timecourse and polysome gradient timecourse with accession numbers GSE299140 and GSE299019, respectively. The results are presented in Figure 6 and Files S6 and S7.

**Figure 6.**
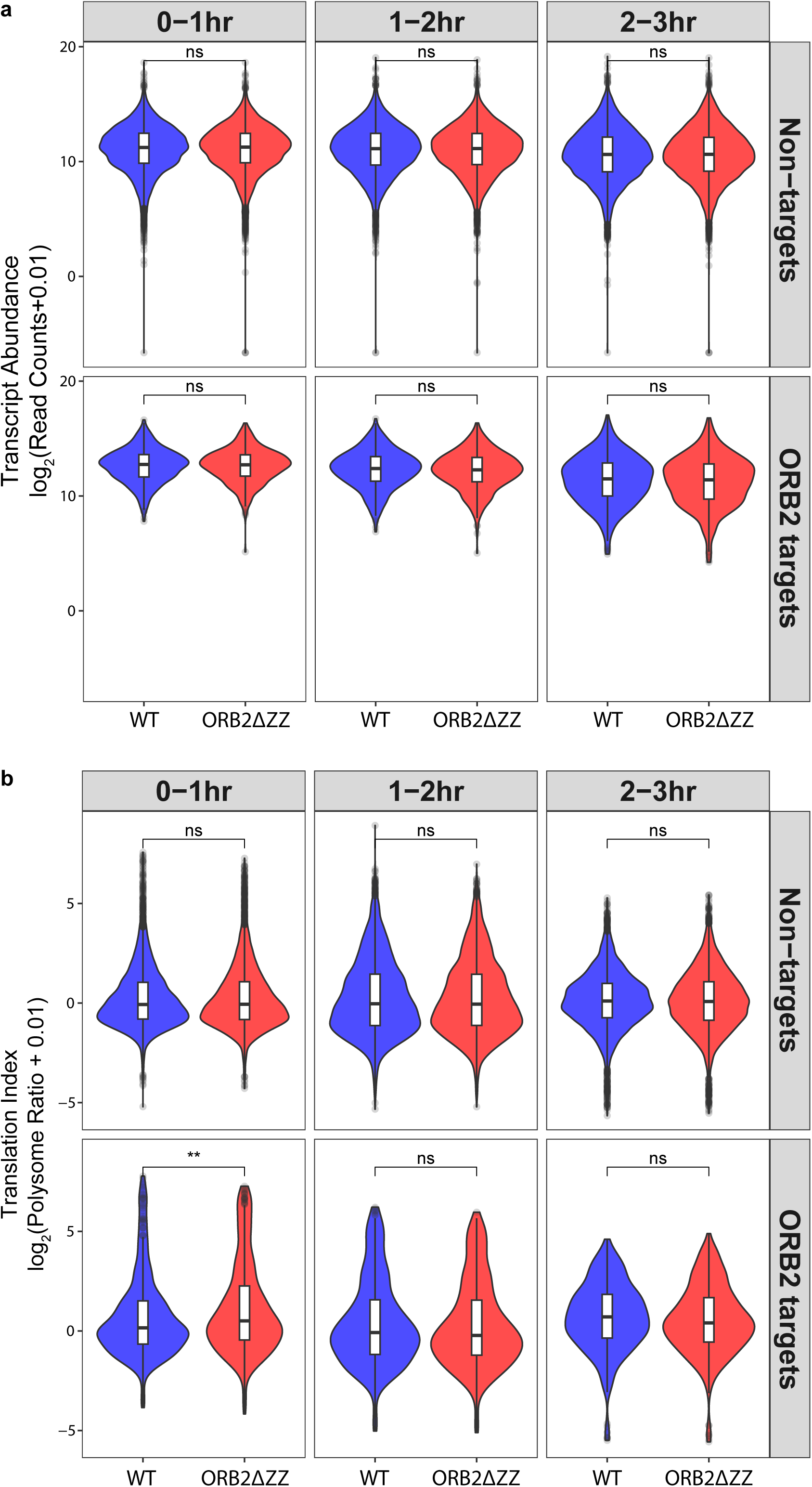
Deletion of the ORB2 ZZ domain results in derepression of target RNAs in the early embryo. (a) Violin and boxplots comparing target and non-target abundance as determined by RNA-Seq in 0-1, 1-2 and 2-3 hour old embryos laid by *w^1118^* and *orb2^ΔZZ^* females. No significant difference was detected for either genotype at any of the time-points. (b) Violin plots and boxplots comparing target and non-target translation index as determined by polysome gradients followed by RNA-Seq in 0-1, 1-2 and 2-3 hour old embryos laid by *w^1118^* and *orb2^ΔZZ^*females. ORB2 targets were significantly derepressed in 0-1 hour old *orb2^ΔZZ^* embryos relative to *w*^1118^. Log2(Ratio of mean normalized counts of pool 3/mean normalized counts of pool 1) was calculated for each transcript. Wilcoxon Rank Sum Test: ** = *P* < 0.01.

### Comparisons between the ORB2 target list and published SMG, PUM, BRAT and RIN target lists

The total list of genes detected in this ORB2 RIP-Seq experiment was first compared with the total list of genes present in the published RIP-Chip experiments for SMG, PUM and BRAT (Chen *et al*. 2014; Laver *et al*. 2015; Laver *et al*. 2020a). The overlap between these lists was retained, and the non-overlapping genes (i.e. genes detected in only one of the two experiments) were discarded. From the remaining genes, ORB2 targets were compared with the targets of SMG, PUM, and BRAT. The translation level, as defined by the polysome RIP-Seq ratio measured in 0-1h embryos (see above), was compared for the overlap between the ORB target list and the other list of interest (i.e. ORB2 + SMG co-targets, ORB2 + PUM co-targets, and ORB2 + BRAT co-targets), and separately for the ORB2-only targets, the SMG- PUM-, and BRAT-only targets, and non targets.

## RESULTS

### ORB2 binds to over 450 targets in the early Drosophila embryo

To identify transcripts that are bound by ORB2B (hereafter referred to as ORB2), we used our previously produced Fab against an antigen N-terminal to the RRMs (Figure 1a) (Laver *et al*. 2012; Na *et al*. 2016) to perform RNA-coimmunoprecipitation (RIP) of endogenously expressed ORB2 from embryo extracts prepared from 0-3 hour old embryos produced by *w^1118^*females. We enriched for mRNA in the sample by rRNA depletion, generated randomly primed cDNA libraries from the resulting samples, and identified co-purifying RNAs by next-generation RNA sequencing (see Materials and Methods). As a negative control, we performed parallel RIPs using a control synthetic antibody, C1, as described previously (Laver *et al*. 2015; Laver *et al*. 2020b; Haugen *et al*. 2024). Input samples from the same extracts were also sequenced. We performed three biological replicates for each of these three conditions, for a total of nine cDNA libraries.

To identify ORB2-bound target genes, we mapped the sequence reads to the Drosophila transcriptome using Salmon read alignment software, and analyzed the results with the DESeq2 software package as described in Materials and Methods (Love *et al*. 2014; Patro *et al*. 2017). We defined an ORB2 target as a gene whose transcripts had fold enrichment >1.5 in the anti-ORB2 RIP-seq compared to input and a ratio of ORB2/Control > 1.5, with an FDR adjusted *P*-value < 0.05; using these criteria, we identified 462 genes whose transcripts were bound by ORB2 (Figure 1b and File S1). ORB2’s target transcripts were significantly less abundant than co-expressed non-target transcripts (Figure 1c). RIP-RT-qPCR verified several of the target and non-target RNAs (Figure 1d).

While ours is the first identification of ORB2 targets in embryos, there are previously published target lists from other tissues or cells. For example, ORB2-bound transcripts were identified in a CLIP experiment derived from ORB2 overexpression in S2 cells (Stepien *et al*. 2016). Of the 462 ORB2 targets on our list, 298 were identified in their CLIP-seq background set of which they identified 113 as targets (Fisher’s exact *P* = 6.52x10^-7^, Odds Ratio = 1.88). We also examined the overlap of our targets with a list of ORB2-associated transcripts obtained by performing RIP-seq on Drosophila head extracts (Kozlov *et al*. 2023). 336 genes in our target gene list overlapped with theirs (Fisher’s exact *P* < 10^-15^, Odds Ratio = 5.00).

The strong overlap between our ORB2 targets and previously reported targets from other cell types/tissues verifies that our list is enriched for true ORB2 interactors.

### ORB2 targets are translationally repressed, deadenylated and degraded during the MZT

To assess the post-transcriptional fate of ORB2 target transcripts in the early Drosophila embryo, we utilized three previously published datasets that scored each transcript’s translational status, poly(A) tail length, and abundance during the MZT.

Two studies measured Translation Efficiency (TE), calculated from ribosome profiling, during the MZT (Dunn *et al*. 2013; Eichhorn *et al*. 2016). At all three timepoints during the MZT that were studied, ORB2 targets had a significantly lower TE compared to co-expressed, non-target transcripts (Figure 2a-c). A third study measured Translation Index (TI), calculated as the ratio of that transcript’s abundance in polysome fractions of a sucrose gradient compared to its abundance in non-polysomal fractions (Chen *et al*. 2014). ORB2 targets had a significantly lower TI than co-expressed non-target transcripts (Figure 2d).

We next assessed changes in poly(A) tail length and abundance for ORB2 targets between the 0-1 hour timepoint and the 2-3 hour timepoint, which had been measured in parallel to TE (Eichhorn *et al*. 2016). Decrease in abundance and poly(A) tail length were significantly greater for ORB2 targets than for co-expressed, non-target transcripts (Figure 2e,f). We also examined previously published lists of transcripts that are degraded during the MZT (Table 1). ORB2 targets overlapped significantly with transcripts identified to be unstable in activated eggs (Tadros *et al*. 2007) (Fisher’s Exact *P* = 2.2 x 10^-16^, odds ratio 3.85). In addition, ORB2 targets were significantly enriched for genes that are degraded during the MZT (Thomsen *et al*. 2010): (Class III and V; respectively, Fisher’s Exact *P* = 0.02, odds ratio 1.54 and Fisher’s Exact *P* = 2.2 x 10^-16^, odds ratio 2.54) and significantly depleted for transcripts that are stable (Class I; Fisher’s Exact *P* = 2.2 x 10^-16^, odds ratio 0.23).

We conclude that ORB2’s target mRNAs are enriched for transcripts that are translationally repressed, deadenylated and degraded during the MZT.

### ORB2 targets are enriched for rare codons

It has previously been reported that ORB2 binds to and upregulates the stability and translation of rare-codon-enriched mRNAs in neurons in *Drosophila* (Stewart *et al*. 2024). We, therefore, asked whether ORB2’s embryonic targets are enriched for rare codons. Two different algorithms – MELP (Supek and Vlahovicek 2005) and CAI (Puigbo *et al*. 2008) – found significant enrichment of rare codons in ORB2 targets relative to non-target, co expressed mRNAs (Figure 2g,h and File S2). Why the post-transcriptional outcome in embryos is the opposite of that in neurons will be considered in the Discussion.

### ORB2 targets are enriched for roles in gamete generation, gene regulation, and development

We next asked whether ORB2’s targets were enriched for particular molecular, cellular or biological functions by performing Gene Ontology (GO) term analysis using the gProfiler2 tool (Gene Ontology *et al*. 2023; Kolberg *et al*. 2023) (Tables 2-4).

The top GO terms under the Molecular Function category (Table 2) were dominated by molecular binding functions; ‘mRNA 3ʹUTR binding’, and ‘mRNA regulatory element binding’ were among the top ten results. Genes belonging to these terms included *orb2* itself as well as *patronin, piwi,* and *bru1*. Other top terms included ‘translation regulator activity’ and ‘translation repressor activity’; genes belonging to these terms included *orb2* itself, *orb, pum, fmr1, sxl, heph,* and *eIF4E-HP*. In the Biological Process category (Table 3), ORB2 targets were enriched for terms related to regulation and development, such as ‘anatomical structure development’, ‘cell differentiation’, and ‘regulation of metabolic process’. ORB2 targets were also enriched for terms related to gametogenesis, including ‘gamete generation’, ‘oogenesis’, and ‘germ cell development’. Genes related to these terms included *pum, rbfox1, piwi, heph, bru1, sxl,* and *mei-P26*. Enrichment for GO terms related to gametogenesis in ORB2 targets is consistent with previously reported functions for ORB2 in, for example, regulation of spermatogenesis (Xu *et al*. 2012; Gilmutdinov *et al*. 2021). The Cellular Component category (Table 4) was dominated by terms that included ‘ribonucleoprotein granule’, ‘P granule’, ‘pole plasm’, and ‘germ plasm’. Genes related to these terms included *larp, tud, osk, tapas, ago1,* and *dcr-2*.

Many of the genes listed under enriched GO terms are previously identified ORB2 targets (e.g., *orb2* itself, *ago1*, *bru1, dcr-2*, *eIF4E-HP*, *fmr1*, *heph*, *orb, pum, rbfox1, unr*) (Mastushita-Sakai *et al*. 2010; Xu *et al*. 2012; Stepien *et al*. 2016; Kozlov *et al*. 2021) while others are previously unidentified players in the same enriched function or process (e.g., *imp* and *piwi*). Enrichment for mRNAs encoding known post-transcriptional regulators suggests that ORB2 may play a key role in the cross-regulatory network that controls the MZT, a function that is supported by our subsequent analyses reported below. The fact that ORB2 binds to and potentially regulates mRNAs that encode other post-transcriptional regulators is also consistent with a previous study in S2 cells showing that RNP complexes tend to be enriched for mRNAs encoding post-transcriptional regulators (Stoiber *et al*. 2015).

### ORB2 target transcripts are enriched for a U-rich motif

CPEB-family proteins are known to bind to U-rich elements, typically located in the 3ʹUTR of their mRNA targets (Hake and Richter 1994; de Moor and Richter 1999; Stepien *et al*. 2016). To predict *cis*-elements recognized by ORB2 from our *in vivo* RIP-seq results, we used our algorithm, Number of Accessible Target Sites (#ATS), which performs *de novo* motif discovery given a target set and a co-expressed non-target set of transcripts (Li *et al*. 2010) as described in the Material and Methods. Using our input RNA-seq data, we identified the most abundant transcript isoform from each expressed gene in the early embryo and then ran #ATS analysis on the full-length transcripts or the 3ʹUTR sequences of ORB2 targets against non-targets. Whereas no motif was produced from the analysis of full-length transcripts, the #ATS program identified a list of 6-mers enriched in the 3ʹUTR of ORB2 target sequences relative to non-targets; these 6-mers were then combined into a consensus motif (Figure 3a, File S3), which will henceforth be referred to as an ORB2-recognition element (ORE: NUUGUWU (where W = A or U). We also calculated an ORE ‘score’ for each transcript by summing the accessibilities of each predicted ORE in that transcript (see Materials and Methods). ORB2 targets had significantly higher ORE scores than co-expressed non-targets (Figure 3b, File S3).

This ORE is similar to published binding motifs: UUUU(A)_1-3_U and UUUUGU identified in S2 cells (Stepien *et al*. 2016); UUUURU (where R = A or G) in brain extracts (Mastushita-Sakai *et al*. 2010); and KUUUKKK (where K = G or U) identified by an *in vitro* binding assay (Ray *et al*. 2013). We calculated ‘KUUUKKK’ scores for each transcript and found that the ORB2 targets we identified in the embryo have significantly higher scores than co-expressed non-targets (Figure S3b, File S3).

The *de novo* discovery of the U-rich ORE enriched in ORB2 target RNAs in the early embryo and its similarity to ORB2 motifs previously identified using a variety of methods supports the conclusion that our list of targets is enriched for transcripts directly bound by ORB2.

### The ORE directs ORB2 binding to and repression of a luciferase reporter mRNA in S2 cells

To validate the biological activity of the ORE identified by #ATS, we used a luciferase reporter assay in Drosophila S2 cells to measure ORB2 binding and ORB2-mediated regulation. In the assay, firefly luciferase (FLuc) reporters harboured six copies in their 3ʹUTR of either a representative ORE (UUUUGUU) or a mutated ORE (UUAUCUC) (the reporter is schematized in Figure 3c). Each of these reporters was individually expressed in S2 cells together with either a control protein (GFP), full-length ORB2 protein, or an ORB2 protein lacking its RBD (ORB2ΔRRM). An unregulated Renilla luciferase (RLuc) reporter was co-transfected with all samples, allowing us to normalize FLuc mRNA and protein levels.

ORB2 RIP followed by RT-qPCR confirmed that the reporters carrying the wild-type OREs, but not the mutant OREs, were enriched in ORB2 IPs, compared to IPs of the ORB2ΔRRM protein (Figure 3d). Furthermore, significant ORB2-mediated repression of the reporter was seen when ORB2 protein was expressed together with a reporter construct carrying wild-type OREs (Figure 3e). When either the OREs were mutated or the RRMs were deleted, no repression was observed (Figure 3e) despite the fact that ORB2ΔRRM protein was expressed at higher levels than full-length ORB2 (Figure S4).

We also assessed whether the firefly luciferase open reading frame used in the reporters might be enriched for *Drosophila melanogaster* rare codons and found that this is indeed the case (Figure S5). Our data do not address whether rare codon-enrichment per se contributes to binding and/or repression; this would require the use of a codon-optimized version of luciferase, which is beyond the scope of this study.

Our data do, however, clearly show that ORB2 binds to OREs in an RBD-dependent manner and that this binding is sufficient to confer repression on the reporter RNA. The magnitude of repression we observed is similar to that previously reported for ORB2B in similar reporter assays in S2 cells (Stepien *et al*. 2016).

### The C-terminal Zinc-binding (‘ZZ’) domain of ORB2/hCPEB2 but not ORB/hCPEB1 represses a luciferase reporter mRNA in S2 cells

To identify domains of ORB2 necessary and sufficient for repression we used a luciferase reporter tethering assay in S2 cells. We transfected a plasmid construct that would express a protein containing the BIV-Tat RNA-binding domain fused to all or part of the ORB2 protein. This fusion protein is recruited to six tandem repeats of the trans-activating response (TAR) element (to which BIV-Tat binds) located in the 3ʹUTR of a co-transfected firefly luciferase reporter (schematised in Figure 4a) (Wakiyama *et al*. 2012; Laver *et al*. 2020a). Tethering of full-length ORB2B protein produced a significant, two-fold repression relative to a tethered GFP control (Figure 4b); the extent of repression was consistent with previously published data for ORB2 in S2 cells (Stepien *et al*. 2016). Tethering of ORB2 protein did not affect steady-state reporter RNA levels (Figure 4c).

To map regions of ORB2 that mediate repression, we fused the BIV-Tat domain to fragments of the ORB2 protein and assessed whether any individual region was capable of repressing luciferase expression. We divided the ORB2 protein into five regions: IDR1, the polyQ region, IDR2, the RBD, and the ZZ domain (Figures 1a and 4b,c). There was no significant effect on the expression of luciferase after tethering of IDR1, polyQ, IDR1+polyQ, IDR2 or RBD (Figure 4b and see Figure S6A for western blot of expression of each fragment). Only the ZZ domain conferred strong (∼three-fold) repression (Figure 4b) despite being expressed at a comparatively low level (Figure S6a). To determine whether the ZZ domain was also necessary for repression, we tethered an ORB2 protein construct lacking the ZZ domain (ORB2ΔZZ) to the luciferase reporter and found that it was unable to repress luciferase expression, despite ORB2ΔZZ retaining the RBD and being expressed at higher levels than full-length ORB2 (Figure 4b, Figure S6b). Because luciferase mRNA levels were not significantly different from controls in either the ZZ sufficiency or the ZZ necessity experiments (Figure 4c, ZZ and ORB2ΔZZ), ZZ domain mediated repression can be attributed largely to effects on translation rather than RNA stability. We conclude that the ZZ domain of ORB2 is both necessary and sufficient to confer translational repression when tethered to a reporter mRNA.

Finally, to assess possible conservation of function, we also tethered the ZZ domains of ORB, hCPEB1 and hCPEB2. The hCPEB2 ZZ domain conferred significant repression, whereas the ORB ZZ and hCPEB1 ZZ domain were unable to repress (Figure 4e; western blot is shown in Figure S6c). We conclude that the repressive function of the ZZ domain is conserved between ORB2 and its human ortholog, hCPEB2, while it is not conserved in the more distantly related proteins, ORB or hCPEB1. We note that the ZZ domains of hCPEB3 and hCPEB4 are ≥ 90% identical to the hCPEB2 ZZ domain (Figure S7a). This suggests that, although not tested here, the hCPEB3 and hCPEB4 ZZ domains likely also function in translational repression.

### Regions within the ORB2 ZZ domain function in combination to confer repression

To map regions of the ORB2 ZZ domain sufficient for repression, we first compared the sequences of the ZZ domains of ORB, ORB2, hCPEB1 and hCPEB2 (Figures 4d and S7a). There is high level of sequence identity between the ORB2 ZZ domain and that of hCPEB2 (81.3% amino acid identity). In contrast, the ZZ domain of ORB2 and ORB, exhibit only 35.6% sequence identity and hCPEB1 has only 28.5% amino acid identity to ORB2. We then considered the predicted structures of these domains (Figure S8). An NMR structure of the hCPEB1 ZZ domain showed it consists of a rubredoxin (Rd) turn, followed by a beta hairpin, and a short alpha helix (Merkel *et al*. 2013). Predictions of the structures of the ORB, ORB2 and hCPEB2 ZZ domains using Alphafold 3 (Abramson *et al*. 2024) showed a similar arrangement. Indeed, Alphafold3 prediction of the ORB ZZ structure was virtually identical to that of the NMR structure of the hCPEB1 ZZ domain (Figure S8a). Similarly, the predictions of the ORB2 and hCPEB2 ZZ structures were very similar to one another (Figure S8a). Finally, an overlap of the ORB and ORB2 ZZ domain structures showed that the only difference was the length of the Rd turn (Figure S8b).

To map regions within the ORB2 ZZ domain sufficient for repression we assessed whether or not chimeric versions that included one or more of the Rd turn (‘A’, Q640 to P656), the beta hairpin (‘B’, F657-Y668), the alpha helix (‘C’, C669 to Y684) and/or the C-terminal region (‘D’, H685 to C-terminus) regions of the ORB or ORB2 ZZ domain functioned to repress firefly luciferase expression using the tethering assay.

As expected, the full-length ORB2 ZZ domain repressed whereas the ORB ZZ domain did not. Of the 14 chimeras tested, three resulted in significant repression relative to the GFP control (Figure 4f, western blots are shown in Figure S7b,c): ORB2 ZZ-A^1^B^2^C^2^D^2^, ORB ZZ-A^2^B^1^C^2^D^1^ and ORB ZZ-A^1^B^1^C^2^D^2^ (where the superscript ‘2’ refers to ORB2-derived sequences and superscript ‘1’ refers to ORB-derived sequences). The region shared by all of these mosaics is ORB2 region C (denoted C^2^), suggesting that it is a major contributor to the repressive function of the ORB2 ZZ domain. However, ORB2 region C alone is not sufficient to confer repression but needs to be combined with another ORB2 region (either A^2^, B^2^ or D^2^) for repression.

### ORB2 interacts with translational repressors and the 43S translation preinitiation complex in early embryos

We next turned to proteomic strategies to identify protein partners of ORB2 in early embryos. We immunoprecipitated endogenous ORB2 from extracts prepared from embryos produced by *w^1118^* females, both in the presence and absence of RNase A, in biological quadruplicates, using the same synthetic antibody used above to identify ORB2 target RNAs. Co-purifying proteins were identified by mass spectrometry followed by Significance Analysis of INTeractome (SAINT), which assigns confidence scores to protein-protein interactions based on comparisons between the Gaussian curves produced by the control IP data and the bait IP data (Choi *et al*. 2012). SAINT was performed with stringent parameters, where for each protein, the lowest control and the highest bait were dropped prior to statistical analysis, which helped ensure that only high-confidence interactors would be identified. We defined a protein interactor of ORB2 as any protein that was significantly enriched in ORB2 immunoprecipitates compared to parallel purifications using our negative control antibody, C1, with a SAINT (AvgP) score ≥ 0.95 and a BFDR < 0.01 (File S4).

We identified 106 interactors in the presence of RNase A (Figures 5a and S10a), representing RNA-independent interactions, and 212 interactors of ORB2 in the absence of RNase A (Figure S10c,d); 85 were in common (File S4). The interactors largely fell into two distinct groups: The first consisted of RBPs known to negatively regulate target stability and/or translation, including AGO1, AGO2, PUM, and RBFOX1, as well as the Cup repressive complex (Cup, ME31B, TRAL, BEL) (Gotze *et al*. 2017) (Figure 5a,c File S4). All components of the Cup repressive complex copurified in the presence of RNase (Figure 5a,c), consistent with an RNA-independent interaction with ORB2. The second group consisted of translation-initiation factors, notably the components of the 43S preinitiation complex (PIC): all eIF3 and eIF2 subunits, eIF1 and -1A, and eIF5A; most (13/20) were RNA-independent interactors, including 11/14 detected eIF3 subunits (Figure 5a,d and File S4).

We note that several of ORB2’s protein interactors were also identified as mRNA targets in our RIP-seq experiments (File S5). This result is consistent with an earlier study of RBP mRNA targets and protein interactors carried out in S2 cells, which showed that RNP complexes are enriched for both mRNAs and their encoded proteins involved in RNA binding and post-transcriptional regulation (Stoiber *et al*. 2015).

Co-purification of ORB2 with post-transcriptional negative regulators is consistent with our analyses of the post-transcriptional fate of ORB2 targets during the MZT and our reporter experiments in S2 cells.

### The ORB2 ZZ domain is required for interaction with the 43S PIC and the Cup repressive complex

Our S2-cell tethering assay, described above, showed that the ZZ domain of ORB2 is necessary and sufficient for translational repression; therefore, we assessed whether this domain is important for association with ORB2’s identified binding partners in early embryos. To generate an endogenous deletion of the ZZ domain, we used a scarless dsRed CRISPR method to remove all amino acids from ORB2 between D638 and the C-terminus at C704, inclusive, leaving the endogenous 3ʹUTR intact (see Figure 4d, Materials and Methods, Figures S1, S2) (Gratz *et al*. 2013; Gratz *et al*. 2014). The mutant fly lines resulting from the excision will henceforth be referred to as *orb2^ΔZZ^*. Western blots on whole-fly extract showed that ORB2ΔZZ was expressed at higher levels than full-length ORB2 in the pre-CRISPRed *w^1118^* wild-type control flies (Figure S1d) and a similar result was obtained for 0-3 hour embryos (Figure S9a). We confirmed by RIP-RT-qPCR that deletion of the ZZ domain did not significantly affect ORB2ΔZZ’s ability to bind to several of its RNA targets in embryos (Figure S9b). As assayed by DAPI staining, *orb2^ΔZZ^* females showed no overt oogenesis defects; females outcrossed to the control, host strain used for CRISPR produced embryos that underwent normal syncytial nuclear cycles, cellularization and gastrulation, and exhibited similar hatch rates into first instar larvae as the control (control: 92.5%. n = 200; *orb2^ΔZZ^*: 90.7%, n = 300). We report elsewhere that *orb2^ΔZZ^*mutants have defects in brain development (Hailstock *et al*. 2025) and spermatogenesis (Low *et al*. 2025).

Immunopurification of ORB2ΔZZ protein from 0-3-hour embryos produced by *orb2^ΔZZ^* homozygous females (henceforth referred to as *orb2^ΔZZ^* embryos) followed by mass spectrometry identified 48 high-confidence protein interactors in the presence (Figure 5b, S10b) and 91 in the absence of RNase A (Figure S10f), of which 43 overlapped (File S4). When the union of these interactors (96 proteins) was compared to the union of full-length ORB2 IP-MS interactors (233 proteins), we found that 65 overlapped, including several translational repressors such as PUM and RBFOX1 (Figure 5b, File S4). Strikingly, interactions with all components of the PIC and the Cup repressive complex were lost upon deletion of the ZZ domain (compare Figure 5a and 5b).

We conclude that the ORB2 ZZ domain is required for interaction between ORB2 and the 43S PIC as well as the Cup repressive complex, but that additional repressors retain interaction with ORB2 in the absence of its ZZ domain.

### Deletion of the ZZ domain results in a shift of ORB2 targets onto polysomes in the early embryo

Since our experiments in S2 cells showed that the ZZ domain is necessary for ORB2-mediated translational repression, and our IP-MS results indicated that ORB2 loses interactions with the Cup repressive complex when the ZZ domain is deleted, we next asked whether deletion of the ZZ domain had consequences for post-transcriptional regulation of ORB2’s target mRNAs during the MZT. Notably, since embryos from *orb2^ΔZZ^* females undergo normal development and binding to targets is unaffected by loss of the ZZ domain, any alterations in the post-transcriptional fate of targets are likely to be a direct effect of the domain deletion rather than a secondary consequence of abnormal development.

First, we assessed whether ORB2’s targets, which are enriched for maternal transcripts that are deadenylated and degraded during the MZT (see Figure 2e,f), were stabilized in *orb2^ΔZZ^* mutant embryos. We collected extracts from embryos laid by control or *orb2^ΔZZ^* females at 0-1, 1-2 and 2-3 hours after egg lay and carried out RNA-seq followed by analysis using DEseq2. We compared the expression level of targets and co-expressed non-targets across this time course and found that, at all three stages, there was no significant difference in expression level between control versus ORB2ΔZZ embryos for either target or non-target transcripts (Figure 6a and File S7). These results suggest that, while ORB2 binds to unstable mRNAs, it does not induce their degradation or, alternatively, that ORB2 may direct destabilization of its targets during the MZT in a ZZ domain-independent mechanism.

Second, we assessed whether ORB2’s targets, which are enriched for transcripts with significantly lower TE and TI than co-expressed non-targets (see Figure 2a-d), underwent an increase in polysome association in ORB2ΔZZ embryos. We collected embryo extracts from embryos laid by control or *orb2^ΔZZ^*females at 0-1, 1-2 and 2-3 hours after egg lay, then ran polysome gradients followed by RNA-seq of pooled polysome fractions and pooled ‘free’ (non-polysome-associated) fractions. For each gene, we calculated the ratio of RNA in the polysome fractions to that in the free fractions (see Materials and Methods); if translational repression were relieved upon deletion of the ZZ domain then this ratio would increase (i.e., transcripts would move onto polysomes). We found that, in ORB2ΔZZ embryos, ORB2 targets shifted onto polysomes at the first timepoint (0-1 hour) whereas non-targets did not (Figure 6b and File S7).

These data are consistent with the hypothesis that the ZZ domain of ORB2 mediates translational repression of ORB2 targets early in the MZT, prior to ZGA. We note that we have not directly demonstrated an increase in protein synthesis from these target transcripts in embryos; however our S2 cell experiments showed that the ZZ domain is both necessary and sufficient to repress luciferase protein expression without a change in reporter RNA levels, consistent with the ZZ domain acting in translational repression.

### Co-regulation of ORB2 target transcripts by additional RBPs

Although the shift onto polysomes of ORB2 targets at 0-1 hours in ORB2ΔZZ embryos versus wild type is statistically significant (*P* < 0.006), it is subtle. We, therefore, decided to assess whether ORB2 might function together with other post-transcriptional regulators known to act during the MZT.

First, we asked whether there was a significant overlap of ORB2’s targets with the targets of several RBPs that we had analyzed previously. We found that ORB2 targets were depleted for targets of Rasputin (RIN), an RBP that positively regulates transcript stability and translation during the MZT (Table 5) (Laver *et al*. 2020a), consistent with a role for ORB2 in negative rather than positive regulation of its targets. In contrast, ORB2 targets significantly overlapped those of PUM, BRAT and SMG (Table 5), RBPs that direct mRNA repression/degradation during the MZT (Tadros *et al*. 2007; Chen *et al*. 2014; Laver *et al*. 2015).

**Table 5.** Overlap of ORB2 targets with those bound by SMG, BRAT, PUM and RIN.

| Comparison | Background | Orb2 Only | Other Only | Overlap | Fisher's Exact<br>p-value | Odds Ratio |
| --- | --- | --- | --- | --- | --- | --- |
| Smaug (Chen et al 2012) | 3758 | 254 | 290 | 41 | 0.0001416 | 2.09148 |
| Brat (Laver et al 2015) | 3347 | 228 | 701 | 69 | 0.02181 | 1.40294 |
| Pum (Laver et al 2015) | 3696 | 238 | 352 | 57 | 6.197E-08 | 2.51398 |
| Rin (Laver et al 2020) | 4429 | 365 | 507 | 6 | 2.15E-10 | 0.14363 |

For each of PUM, BRAT and SMG, we subdivided ORB2’s targets into ones bound only by ORB2, ones bound by both ORB2 and that RBP, ones bound only by that RBP, and co-expressed unbound transcripts (see Materials and Methods, File S8). We found that, upon loss of the ZZ domain, ORB2 targets bound by ORB2 but not SMG exhibited a significant shift onto polysomes (*P* = 0.009; Figure 7b ‘ORB2 only’) whereas those co-bound by SMG (Figure 7a), bound by SMG but not ORB2 (Figure 7c) or bound by neither ORB2 nor SMG (Figure 7d) did not significantly shift onto polysomes. This result is consistent with the hypothesis that co-regulation of a subset of ORB2’s targets by SMG maintains repression (i.e., they do not shift onto polysomes when the ZZ domain is deleted) and explains at least in part the subtle albeit significant shift seen when ORB2 targets are aggregated (Figure 6b). Consistent with the SMG analyses, ORB2 targets co-regulated (or not) by BRAT and PUM, exhibited a similar trend but with the ORB2-only targets just missing significance (*P* = 0.062 and 0.073 for the PUM and BRAT comparisons, respectively; Figures S11 and S12).

**Figure 7.**
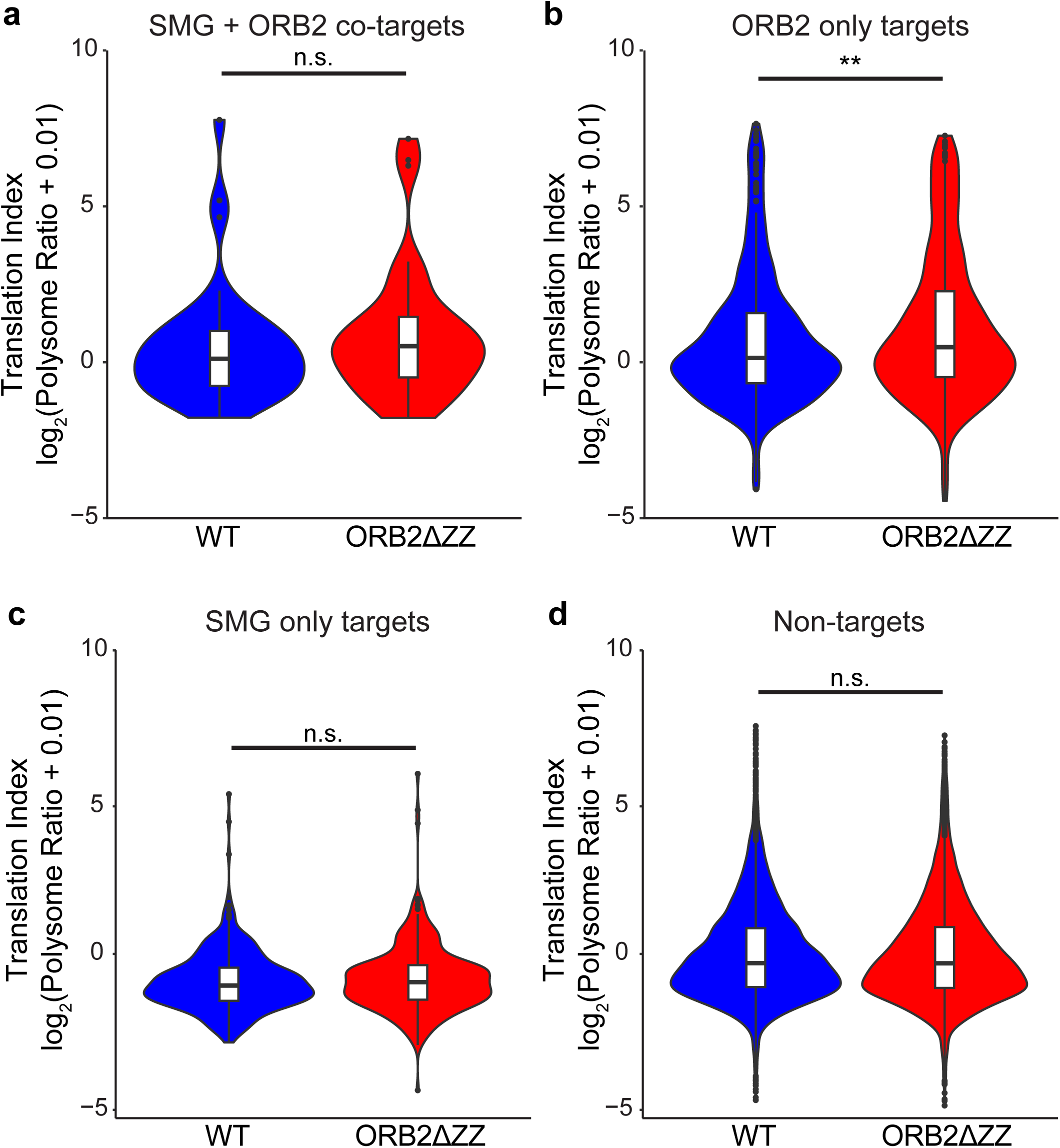
ORB2 targets which are co-bound by SMG are not repressed in the early embryo. (a-d) Violin and boxplots comparing the polysome association of ORB2 and SMG co-targets (a), ORB2 targets (b), SMG targets (c), and non-targets (d), as determined by polysome gradients followed by RNA-Seq in 0-1 hour old embryos laid by *w^1118^* control (WT) and *orb2^ΔZZ^* females. Wilcoxon Rank Sum Test ** = *P* < 0.01.

These data are consistent with the hypothesis that a significant subset of ORB2’s targets are coregulated by additional repressive RBPs, which attenuate the effect of loss of ORB2 repression upon deletion of the ZZ domain.

## DISCUSSION

In this study, we identified hundreds of RNA targets and protein partners of the Drosophila ORB2 protein in the early embryo. ORB2’s targets are enriched for translationally repressed mRNAs that are degraded during the MZT. ORB2-target open reading frames are enriched for rare codons and a U-rich motif in their 3ʹUTR, the latter of which directs binding and repression of a reporter in S2 cells. Repression by ORB2 is mediated by the ZZ domain, a function that is conserved in hCPEB2. Deletion of the ZZ domain from the endogenous ORB2 gene results in loss of physical interaction with the Cup repressive complex and movement of ORB2’s mRNA targets onto polysomes in the early embryo. Subsets of ORB2’s targets are co-bound and likely co-regulated by SMG.

### A conserved role for the ORB2/hCPEB2 ZZ domain in mediating translational repression

Structural analysis of the ZZ domain of hCPEB1 has implicated it in protein-protein interaction (Merkel *et al*. 2013; Stepien *et al*. 2016), consistent with earlier studies on, for example, the ZZ domain of CREB-binding protein (Legge *et al*. 2004). Our identification of multiple proteins that depend *in vivo* on the ZZ domain for interaction with ORB2 are consistent with a major role in recruiting co-regulators to target transcripts. Our data showing that deletion of the ZZ domain does not affect RNA binding are consistent with a previous study (Stepien *et al*. 2016). The fact that, in our S2 cell tethering assay, the ZZ domain is necessary and sufficient for repression permitted us to assess evolutionary conservation of this role as well as to map regions of the ZZ domain that implement this function.

Interestingly, we found that the repressive function of the entire ZZ domain is conserved between ORB2 and hCPEB2 but not with ORB and hCPEB1. Chimeras containing regions from the ORB2 and ORB ZZ domain showed that the α-helix (region C, C669 to H685) of ORB2 can convert the ORB ZZ domain into a repressor when paired with one of the following: ORB2’s Rd turn at the N-terminus (Region A, Q640 to P656), the beta hairpin (Region B, F657-Y668), or the C-terminal beta-sheet region (Region D, H685 to C-terminus). The ZZ domains of hCPEB3 and 4 show ≥ 90% amino acid identity to hCPEB2, with 16/17 residues of region A, 14/15 of region B, 12/13 of region C, and 15/20 of D identical between hCPEB2-4. As such, it is quite likely that the ZZ domains of hCPEB3 and 4 are also capable of repressing target mRNA translation via region C.

That fact that region C of ORB2’s ZZ domain needs one of A, B or D to convert the ORB ZZ domain into a translational repressor is consistent with a model in which A, B or D bind different co-repressors, any of which cooperate with C to mediate repression, or a model in which A, B or D establish a structural context for C to interact with co-repressor(s). Future work aimed at identifying the ORB2 binding partners that interact with each of regions A-D should provide insights into the molecular mechanisms that underlie ZZ domain function.

### Regulation of post-transcriptional processes by ORB2 during the MZT: features of ORB2’s targets and their implications

Three features in ORB2’s target transcripts have implications for post-transcriptional regulation during the MZT: enrichment for OREs, enrichment for rare codons, and enrichment for binding sites for additional RBPs (notably SMG, BRAT and PUM).

First, enrichment for OREs and our evidence from S2 cell reporter assays that ORB2 binds to OREs, which then confers repression on the reporter, are consistent with previous studies of the binding specificity of ORB2 and ORB2B’s role as a repressor (Khan *et al*. 2015; Stepien *et al*. 2016). Notably, since ORB2B is the only isoform expressed during the MZT, we conclude that the *orb2* gene’s function during the MZT is exclusively negative, which contrasts with its role in other tissues. For example, in the nervous system, ORB2A and ORB2B are co-expressed, permitting hetero-oligomer formation and translational upregulation of ORB2 targets (Majumdar *et al*. 2012; Khan *et al*. 2015). Thus, expression specifically of the ORB2B isoform during the MZT is a mechanism to negatively regulate the post-transcriptional fate of its targets.

Second, enrichment of ORB2’s targets for rare codons is consistent with previous observations in neurons, where it has been shown that ORB2 binds rare-codon enriched transcripts (Stewart *et al*. 2024). In contrast to neurons, where the outcome is upregulation of transcript stability and translation, embryonic targets of ORB2 are repressed and degraded. As discussed above, this may be in part due to an absence of ORB2A and a consequence of only having ORB2B present, but it may also be because of additional specific, negative-regulatory co-factors that interact with ORB2 and/or ORB2 targets in embryos, as discussed in the next section. Either way, the observation that ORB2 targets are enriched for rare codons and are downregulated is consistent with the larger literature on the outcome of non-optimal codon usage, including during the MZT of flies and vertebrates (Presnyak *et al*. 2015; Bazzini *et al*. 2016; Iriarte *et al*. 2021).

Third, we have shown that significant subsets of ORB2’s targets are co-bound during the MZT by SMG, BRAT and PUM. Our analyses of the ORB2 targets that move onto polysomes when the ZZ domain is deleted are consistent with the hypothesis that co-regulated targets (e.g., those bound by both ORB2 and SMG) continue to be repressed, likely because SMG directs ongoing repression, whereas ORB2-targets not bound by SMG shift onto polysomes. Whether the latter are co-bound by additional unidentified RBPs is not yet known, but the fact that they move onto polysomes suggests that, even if co-bound by such RBPs, those RBPs are not sufficient for repression, at least not at the level of translation initiation.

### Regulation of post-transcriptional processes by ORB2 during the MZT: the role of repressive co-factors

Our analyses of the post-transcriptional fate of ORB2’s targets during the MZT indicate that they are enriched for transcripts that are translationally repressed and degraded. Consistent with this outcome, multiple co-factors that are known to mediate repression and degradation of mRNAs copurify with ORB2 in early embryos; most of these copurify in the presence of RNase, consistent with an RNA-independent interaction with ORB2. Several of the regulators that interact in an RNA-independent manner require the ZZ domain for interaction, notably the Cup complex components Cup, TRAL, ME31B and BEL.

We have shown that the ZZ domain is necessary and sufficient for translational repression of reporters in S2 cells and deletion of the ZZ domain results in a shift of ORB2’s *in vivo* targets onto polysomes in 0-1-hour embryos. That the Cup complex is expressed in both S2 cells and embryos, that ZZ domain deletion results in loss of Cup complex interaction with ORB2 in embryos, and that ORB2’s targets in embryos move onto polysomes when the ZZ domain is deleted from endogenous ORB2, are all consistent with the hypothesis that ORB2-dependent repression is mediated by the Cup complex. The Cup complex prevents interaction of eIF4G with eIF4E, thus abrogating translation initiation (Wilhelm *et al*. 2003; Nakamura *et al*. 2004; Nelson *et al*. 2004a); consistent with this role, we note that, while eIF4E was present in our ORB2 IP-MS data, eIF4G was absent. Previous studies have shown that the Cup complex is brought to mRNAs by BRU or Squid (SQD) in oocytes and by SMG in embryos (Wilhelm *et al*. 2003; Nakamura *et al*. 2004; Nelson *et al*. 2004a; Clouse *et al*. 2008). ORB2, thus, represents a second RBP that acts in embryos to recruit this complex for repression, in this case early in the MZT. A caveat is that we have not demonstrated that the targets that shift onto polysomes are translated; post-initiation mechanisms may act to prevent protein synthesis.

The Cup complex has been shown to be cleared by the CTLH E3 ligase part-way through the MZT (Cao *et al*. 2020; Zavortink *et al*. 2020), consistent with the ZZ domain’s repressive role in the early but not late MZT. However, ORB2’s targets are repressed at both 0-1 hours (when the Cup complex is present) and at 2-3 hours (after the Cup complex has been cleared); furthermore, ORB2’s targets are deadenylated and degraded between 0-1 and 2-3 hours, but deletion of the ZZ domain has no effect on their clearance. These data suggest that additional regulators may function either independent of or together with ORB2 to direct these processes. SMG is an example of a co-regulator present throughout the MZT that acts through the Cup complex for repression and through the CCR4-NOT deadenylase to direct degradation (Nelson *et al*. 2004b; Semotok *et al*. 2005). SMG’s co-purification with ORB2 in our IP-MS experiments is RNA-dependent; thus, SMG’s binding and regulation of a subset of ORB2-targets may be independent of physical interaction with ORB2. PUM, FMR1 and RBFOX1 are also present throughout the MZT (Casas-Vila *et al*. 2017; Cao *et al*. 2020) and we have shown here that their interaction with ORB2 is both RNA-independent and ZZ-domain independent, making them candidates for implementing these roles together with ORB2 but independent of its ZZ domain.

### The mystery of PIC interaction

Unexpectedly, we identified the 43S PIC as a major interactor of ORB2 via its ZZ domain. An essential component of the PIC is the eIF3 complex (its subunits are encoded by 14 genes in *Drosophila*), which is a highly regulated component of translation initiation (Harris *et al*. 2006; Chiu *et al*. 2010; Cate 2017). Several of the mammalian eIF3 subunits have been shown to have RNA-binding ability (Lee *et al*. 2015; Lee *et al*. 2016) and to mediate distinct regulatory outcomes (Pestova *et al*. 1998; Pestova and Hellen 1999; Hui *et al*. 2003; Hui *et al*. 2005; Martineau *et al*. 2008; Lee *et al*. 2010; Martineau *et al*. 2014).

At this time we do not have data that provide insights into the functional consequences of ORB2’s interaction with the PIC. ORB2’s interaction with the PIC’s eIF3 subunits is RNA-independent, consistent with direct interaction with one or more of these subunits. We considered one possibile mechanism where ORB2 may sequester the PIC, thus limiting its availability and reducing translation overall. However, in such a model, one would expect a global increase in translation upon loss of PIC interaction rather than what we observed in *orb2^ΔZZ^* embryos, which is an increase in polysome association of ORB2’s targets but no change in non-targets. An alternative hypothesis is that ORB2-dependent binding to the PIC on target transcripts has a negative effect on translation, either in addition to or in concert with the Cup repressive complex. Indeed, a speculative model might be that ORB2 binding to the 3ʹUTR and its interaction with PIC at the 5ʹUTR might repress translation initiation by interfering with PIC function or enhancing the function of repressive elements of the eIF3 complex. The mechanistic and functional consequences of ORB2-PIC interaction will be a fruitful area for future study.

## Data Availability

Drosophila strains are available upon request. Raw data for RIP-seq, RNA-seq and IP-MS are available at GEO (GSE299661 for RIP-seq, GSE299140 for RNA-seq time-course, [GSE299019](https://www.ncbi.nlm.nih.gov/geo/query/acc.cgi?acc=GSE299109) for polysome gradient seq) and at MassIVE ([MSV000098275](https://massive.ucsd.edu/ProteoSAFe/private-dataset.jsp?task=843e675042b14ed1a140977c810c427e) for IP-MS). The authors affirm that all of the other data necessary for confirming the conclusions of the article are present within the article, figures, tables and supplementary files.

## Acknowledgements

We thank Kun Nie for assisting with the *de novo* motif discovery using #ATS, Dr. Dorothy Lerit and Dr. Donald Fox for feedback on draft and preprint versions of this manuscript, respectively.

## Supplemental Figure Legends

**Figure S1. CRISPR Cas9-mediated endogenous deletion of the ORB2 ZZ domain was used to generate *orb2*^ΔZZ^ flies.** (a) Diagram depicting the *orb2* endogenous locus, located at 66E4-66E5 on 3L, the corresponding mRNAs that are encoded, and the targeted deletion. (b) Agarose gel of the PCR amplification of the region surrounding the CRISPR targeted deletion of the F1 generation post-injection. Genotypes indicate lines generated from individual F1 flies. Up PCR indicates PCR amplification using primers designed against the 5ʹ insertion junction; Down PCR indicates PCR amplification using primers designed against the 3ʹ insertion junction. (c) Agarose gel of the PCR amplification of the region surrounding the PiggyBac transposon-mediated excision of the dsRed marker. WT indicates the band representing PCR amplification of the endogenous ORB2 locus containing an intact ZZ domain; Excised indicates the band representing amplification of the locus with the ZZ domain deletion without the dsRed marker. Genotypes indicate lines generated from individual flies resulting from the CRISPR injection. (d) Western blot showing expression of endogenous ORB2 and CRISPRed ORB2ΔZZ in heterozygous and homozygous whole flies. ORB2ΔZZ is expressed at higher levels than full-length ORB2 (see heterozygote for comparison). Generation of CRISPR mutants was performed by WellGenetics as described in the Materials and Methods.

**Figure S2. Sequence of the CRISPR Cas9-mediated endogenous deletion of the ORB2 ZZ domain.**

**Figure S3. Replicate-to-replicate comparisons of transcript TPMs from RIP-seq experiments show high correlation between replicates.** (a)-(c) Pairwise comparisons of the three biological replicates performed with the anti-ORB2 E8 Fab. The Pearson correlation coefficient and its corresponding *P*-value were calculated and used to determine the degree of similarity between each pair. (d) Heatmap of the Poisson distance between three biological RIP-seq replicates performed with anti-ORB2 E8 Fab and three biological RIP-seq replicates performed with control C1 Fab. The Poisson distance is calculated by the difference in Poisson distribution between pairs of samples; a larger distance indicates a greater level of dissimilarity between samples.

**Figure S4. Evaluation of RNA Compete motif enrichment and expression levels of ORB2, ORB2ΔRRM, and GFP in S2 cells.** (a) The transcriptome was evaluated for accessible enrichment of the KUUUKKK motif and each transcript was scored based on the number and accessibility of these motif sites, as in Figure 3b. ORB2 targets had significantly higher scores compared to non-targets, as assessed by Wilcoxon rank sum test. *** = *P* < 10^-15^. (b) Each set of three lanes corresponds to the ORE validation experiments in which expression vectors for tagged proteins and reporter mRNA constructs containing various recognition motifs were transfected into S2 cells, described in Figure 3d and 3e.

**Figure S5. The firefly luciferase open reading frame used in the FLuc reporter is enriched for rare codons.** (a) Distribution of CAI scores, calculated for all annotated CDS longer than 80 codons on FlyBase 6.58 by local version of CAIcal with the *Drosophila melanogaster* codon usage table from the Kazusa codon usage database, and the isoform with minimum score (the most enriched for rare codons) is chosen to represent each expressed gene. (b) Distribution of MELP scores, calculated for all annotated CDS longer than 80 codons on FlyBase 6.58 by coRdon with ribosomal proteins as the references for codon optimization, and the isoform with minimum score (the worst expressivity) is chosen to represent each expressed gene. In (a) and (b) the scores for the firefly luciferase open reading frame are indicated by the dashed blue lines.

**Figure S6. Expression of ORB2 full-length protein, portions of ORB2, and the ZZ domains of ORB, ORB2, hPCPEB1 and 2 in S2 cells.** (a) Western blot showing the expression of ORB2 full-length protein and protein portions corresponding to the experiments described in Figure 4b and c. (b) Western blot showing the expression of ORB2B and ORB2BΔZZ to the experiments described in Figure 4b and c. (c) Western blot showing the expression of ORB2 ZZ, hCPEB2 ZZ, ORB ZZ and hCPEB1 ZZ corresponding to the experiments described in Figure 4e.

**Figure S7. Expression of chimeric ORB-ORB2 ZZ domain proteins in S2 cells.** (a) Schematic depicting the amino acid sequence alignment of the ZZ domains of ORB2, ORB, and their human orthologs, hCPEB1, hCPEB2, hCPEB3, and hCPEB4. The arrowhead denotes the junction between the RBD and ZZ domains. (b) Bar graph depicting the expression of the reporter firefly luciferase when ORB ZZ chimeric domains harboring substitutions from the ORB2 ZZ domain, are tethered. The ORB2 ‘A’ and ‘C’ regions are each sufficient to convert the ORB ZZ domain into a repressor. One-way Kruskal-Wallis non-parametric test with Dunn multiple test correction was used to assess significance. (c) Western blot showing the expression of ORB-ORB2 chimeric proteins corresponding to the substitution experiments which assessed transformation of the non-repressing ORB ZZ domain into a translational repressor, as shown in S7b above.

**Figure S8. Structure of ZZ domains of CPEB family members.** (a) The NMR structure of the ZZ domain of human CPEB1 Protein Data Bank accession 2M13 (Merkel *et al*. 2013) is shown, along with the Alphafold 3 (Abramson *et al*. 2024) predicted structures of the ZZ domains of ORB, hCPEB2 and ORB2. The colors in the Alphafold 3 predictions represent the predicted local distance difference test (pLDDT) scores, where higher numbers indicate greater confidence in the prediction. (b) On the left is an overlap of the ORB and ORB2 predicted structures with Region A (the Rd turn), Region B (the beta hairpin), Region C (the alpha helix), and Region D (the beta sheet) of each protein indicated. The boxes on the right depict the most confident structure within each region in two different orientations.

**Figure S9. ORB2ΔZZ retains RNA-binding to its targets in embryos.** (a) Western blot showing the expression of ORB2 and ORB2ΔZZ protein over the first four hours of embryonic development. (b) Bar graph depicting the enrichment of five ORB2 target mRNAs in RIP-RT-qPCR from 0-3 hour old embryos laid by *w*^1118^ and *orb2*^ΔZZ^ females. The ORB2 RIPs used the ORB2 E8 Fab; the control RIP used the C1 Fab. RNA levels were normalized to *RpL32* mRNA. n = 3; error bars are standard deviation. Student’s *t*-test *P-*values were not significant (n.s.; *P* > 0.05)

**Figure S10. IP-MS identifies ORB2 protein interactors.** (a,b,d,e) Scatterplots depicting all proteins identified in the IP-MS. Log_2_(average spectral counts) from the ORB2 IP is plotted against the Log_2_(average spectral counts) from the control IP. (a) IP-MS performed on embryos laid by *w*^1118^ females in the presence of RNase (corresponding to Figure 5a), (b) IP-MS performed on embryos laid by *orb2*^ΔZZ^ females in the presence of RNase (corresponding to Figure 5b), (d) IP-MS performed on embryos laid by *w*^1118^ females in the absence of RNase, and (e) IP-MS performed on embryos laid by *orb2*^ΔZZ^ females in the absence of RNase. (c, d) Scatterplots depicting all proteins identified in the IP-MS performed in the absence of RNase. Enrichment is expressed as the Log_2_ ratio of the average spectral counts detected in the ORB2 IP compared to the average spectral counts detected in the control IP. Protein interactors were identified by SAINT (with a SAINT score ≥ 0.95 and BFDR < 0.01), shown in red.

**Figure S11. ORB2 targets which are co-bound by PUM are not de-repressed in the early embryo.** (a-d) Violin plots and boxplots comparing the translation index of ORB2 and PUM co-targets (a), ORB2 targets (b), PUM targets (c), and non-targets (d), as determined by polysome gradients followed by RNA-Seq in 0-1 hour old embryos laid by *w^1118^* and *orb2^ΔZZ^*females. Wilcoxon Rank Sum Test *P-*values are shown.

**Figure S12. ORB2 targets which are co-bound by BRAT are not de-repressed in the early embryo.** (a-d) Violin plots and boxplots comparing the translation index of ORB2 and BRAT co-targets (a), ORB2 targets (b), BRAT targets (c), and non-targets (d), as determined by polysome gradients followed by RNA-Seq in 0-1 hour old embryos laid by *w^1118^* and *orb^2ΔZZ^* females. Wilcoxon Rank Sum Test *P*-values are shown.

## Supplemental File Legends

**File S1. Results of RIP-seq of ORB2 in 0-3 hour embryos**. See Materials and Methods for details.

**File S2. Results of rare codon enrichment analyses on ORB2 targets in 0-3 hour embryos.** The MELP and CAI were calculated as described in Materials and Methods.

**File S3. Results of *de novo* motif discovery on ORB2 targets in 0-3 hour embryos.** Motif discovery was carried out using #ATS as described (Li *et al*. 2010). ORE scores and ‘KUUUKKK’ scores are listed. See Materials and Methods for details.

**File S4. Results of IP-MS of ORB2 and ORB2ΔZZ in 0-3 hour embryos in the presence or absence of RNase A.** Interactors were identified using SAINT as described (Choi *et al*. 2012). See Materials and Methods for details.

**File S5. Lists of mRNAs bound by ORB2 (RIP-seq) whose protein products were also identified as interactors in ORB2 IP-MS.**

**File S6. Results of RNA-seq time-course in wild-type and ORB2ΔZZ embryos.** See Materials and Methods for details.

**File S7. Results of polysome gradient RNA-seq time-course in wild-type and ORB2ΔZZ embryos.** See Materials and Methods for details.

**File S8. Overlaps of ORB2 targets and previously identified SMG, BRAT and PUM targets.** See Materials and Methods for details.

