## Supplementary figures and images for "The ORB2 RNA-binding protein negatively regulates its target transcripts during the Drosophila maternal-to-zygotic transition via its functionally conserved Zinc-binding ‘ZZ’ domain"

### Figure S1

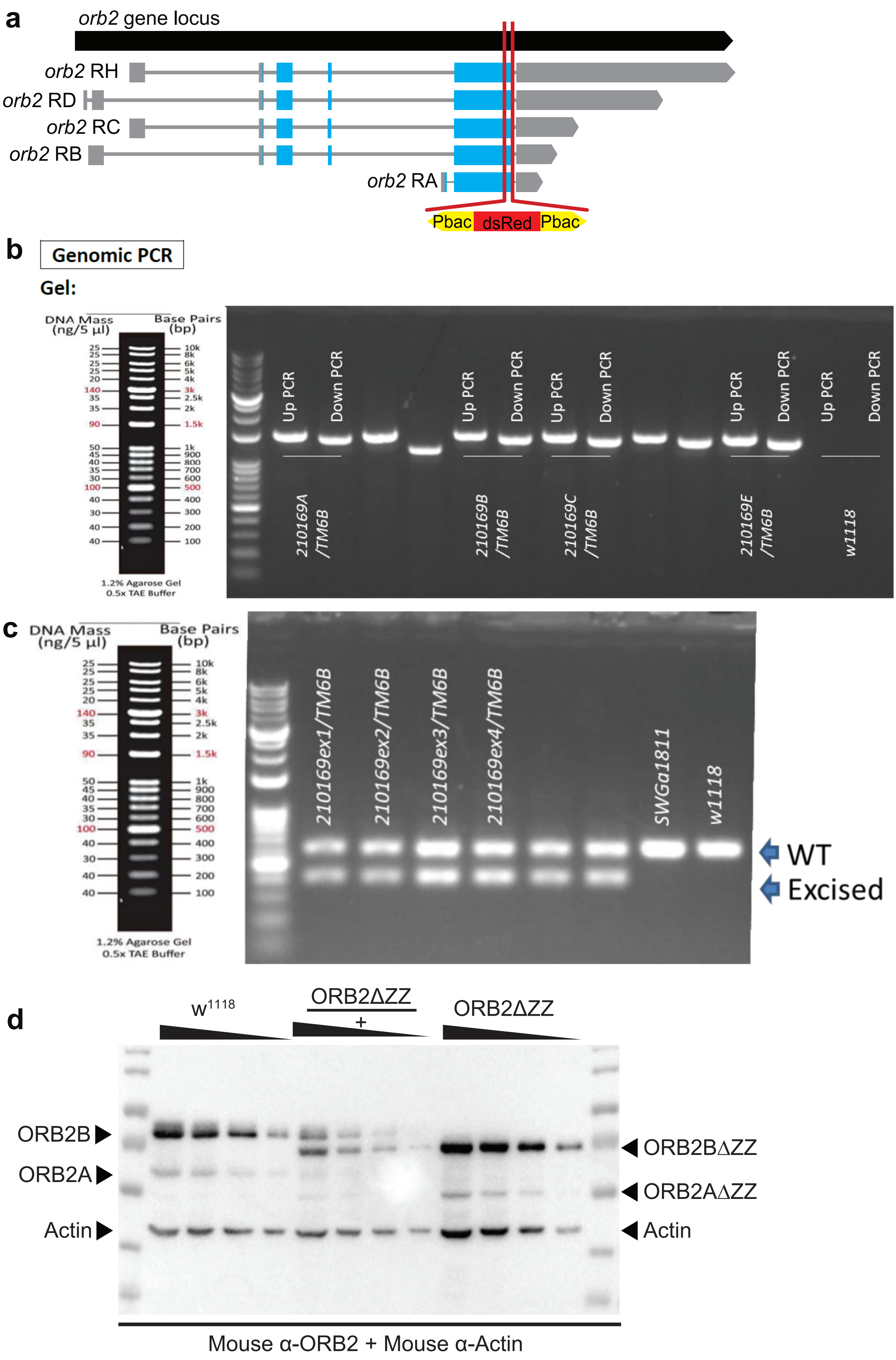

### Figure S3

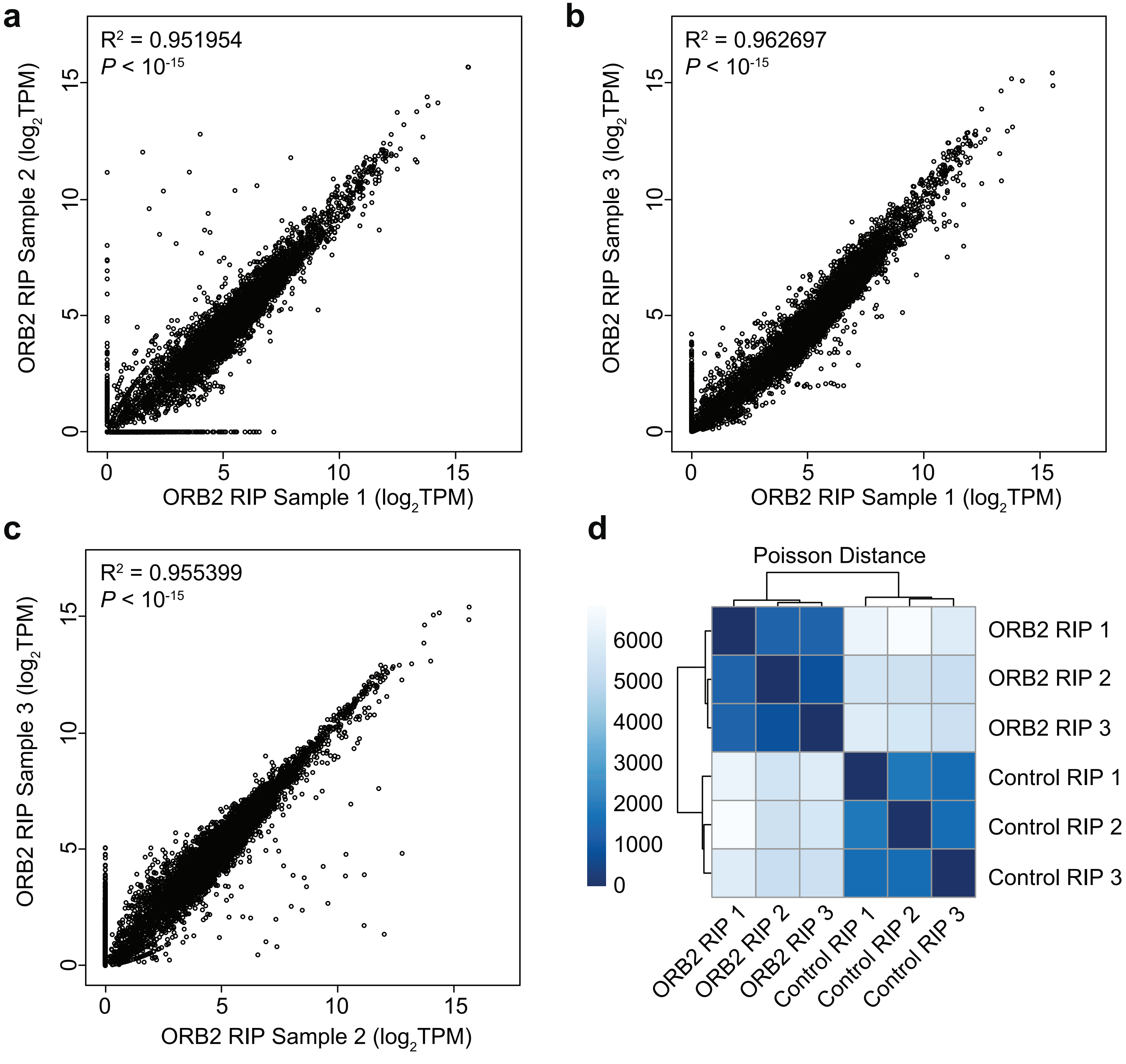

### Figure S4

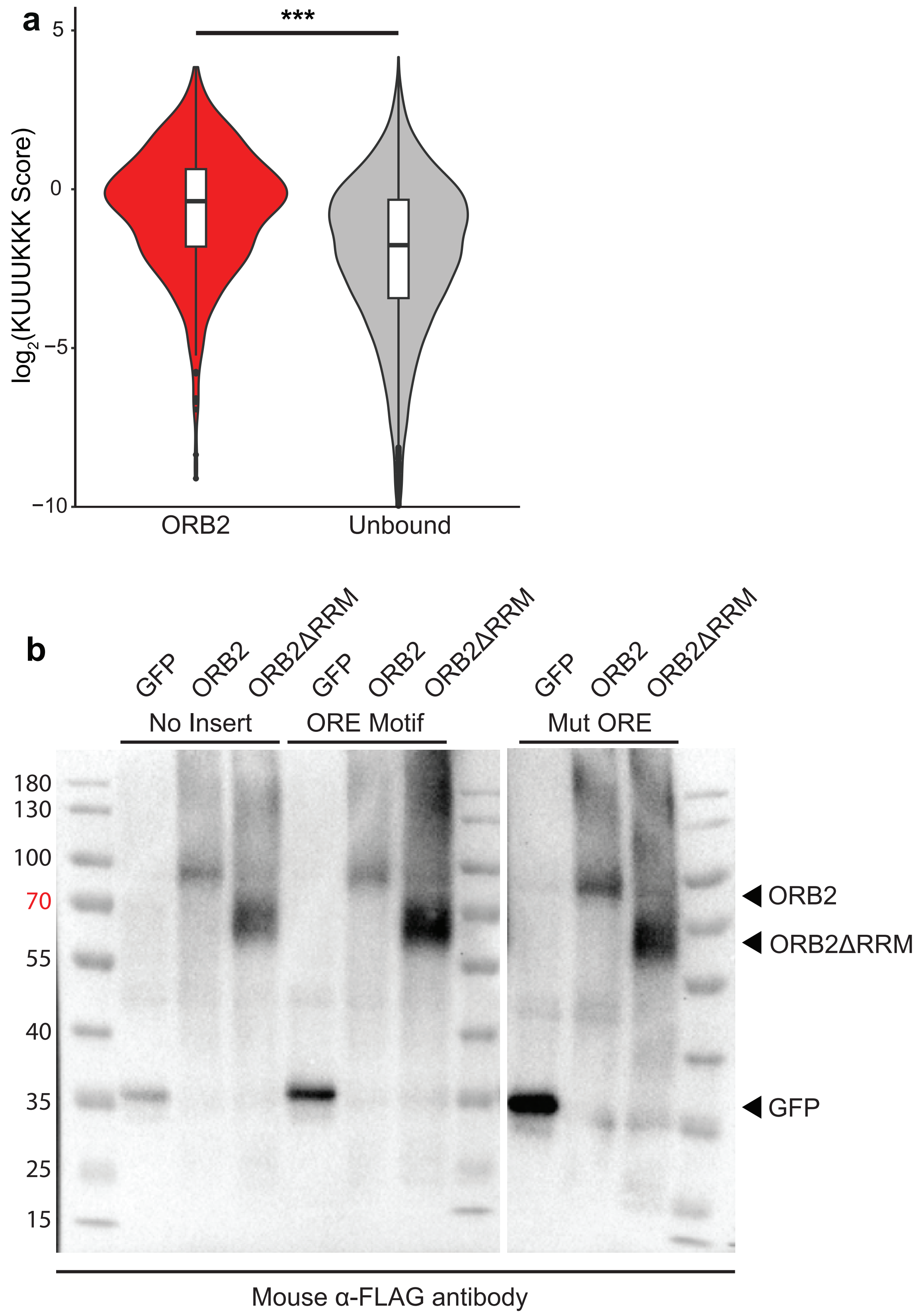

### Figure S5

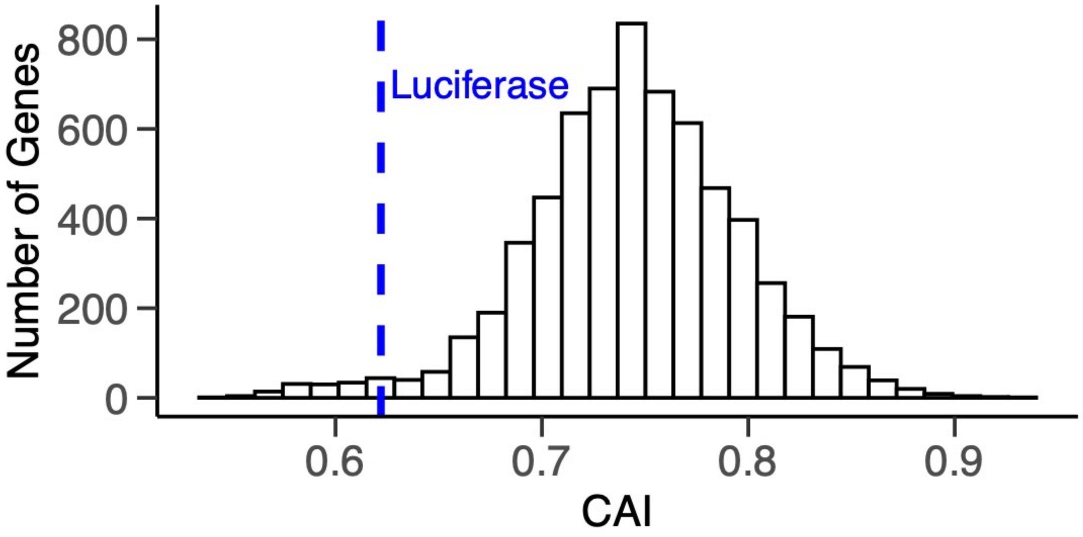

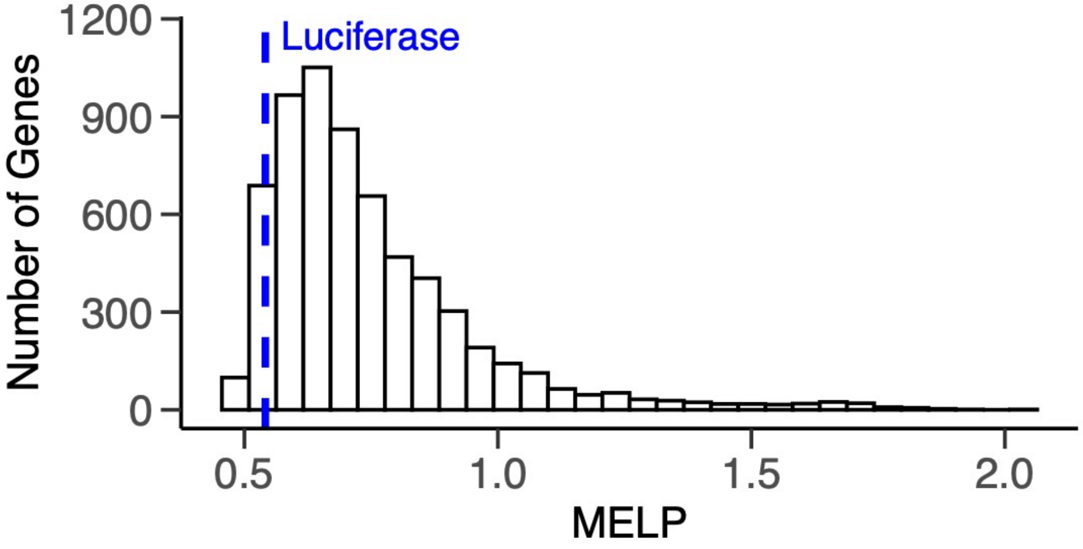

### Figure S6

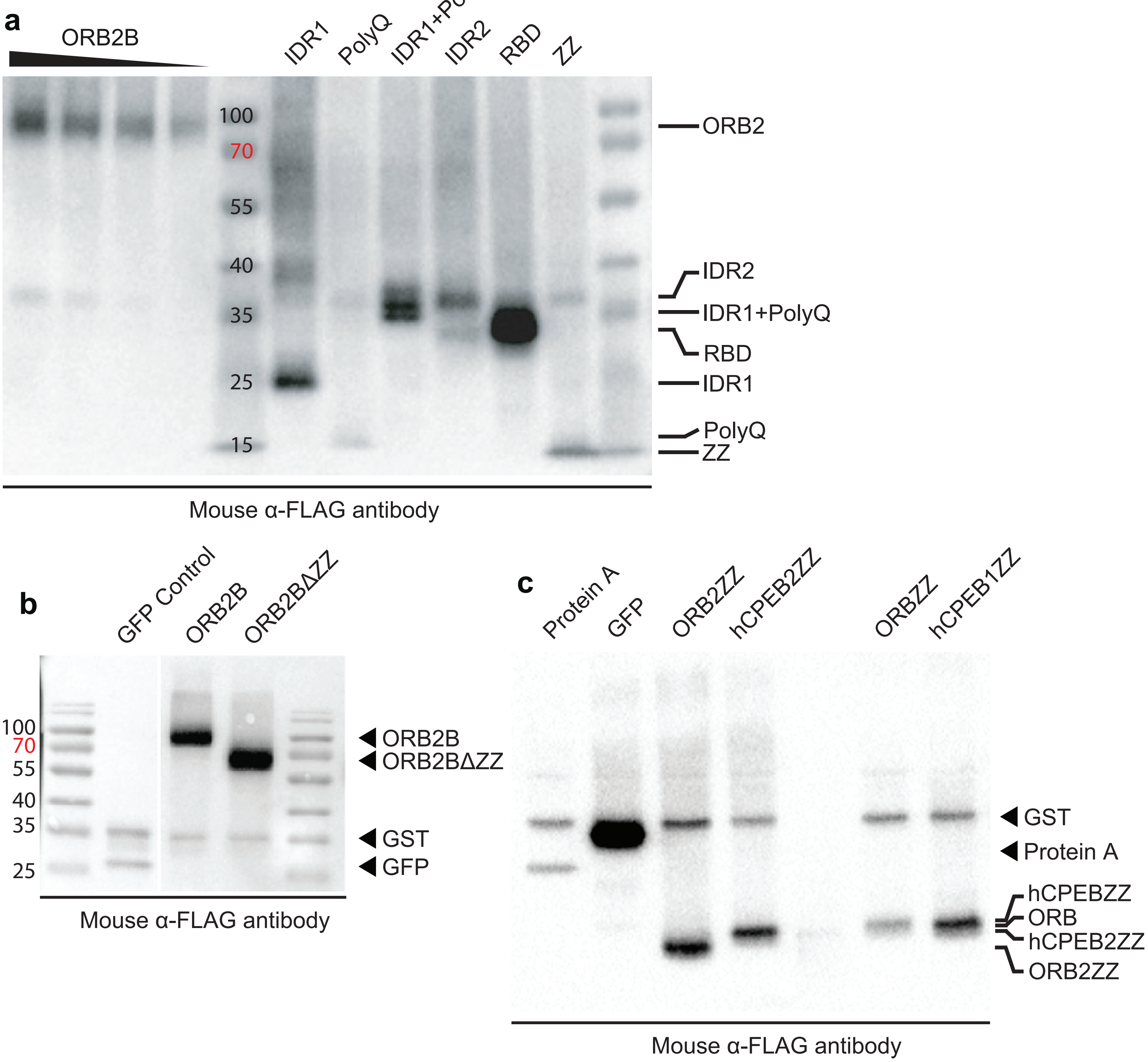

### Figure S7

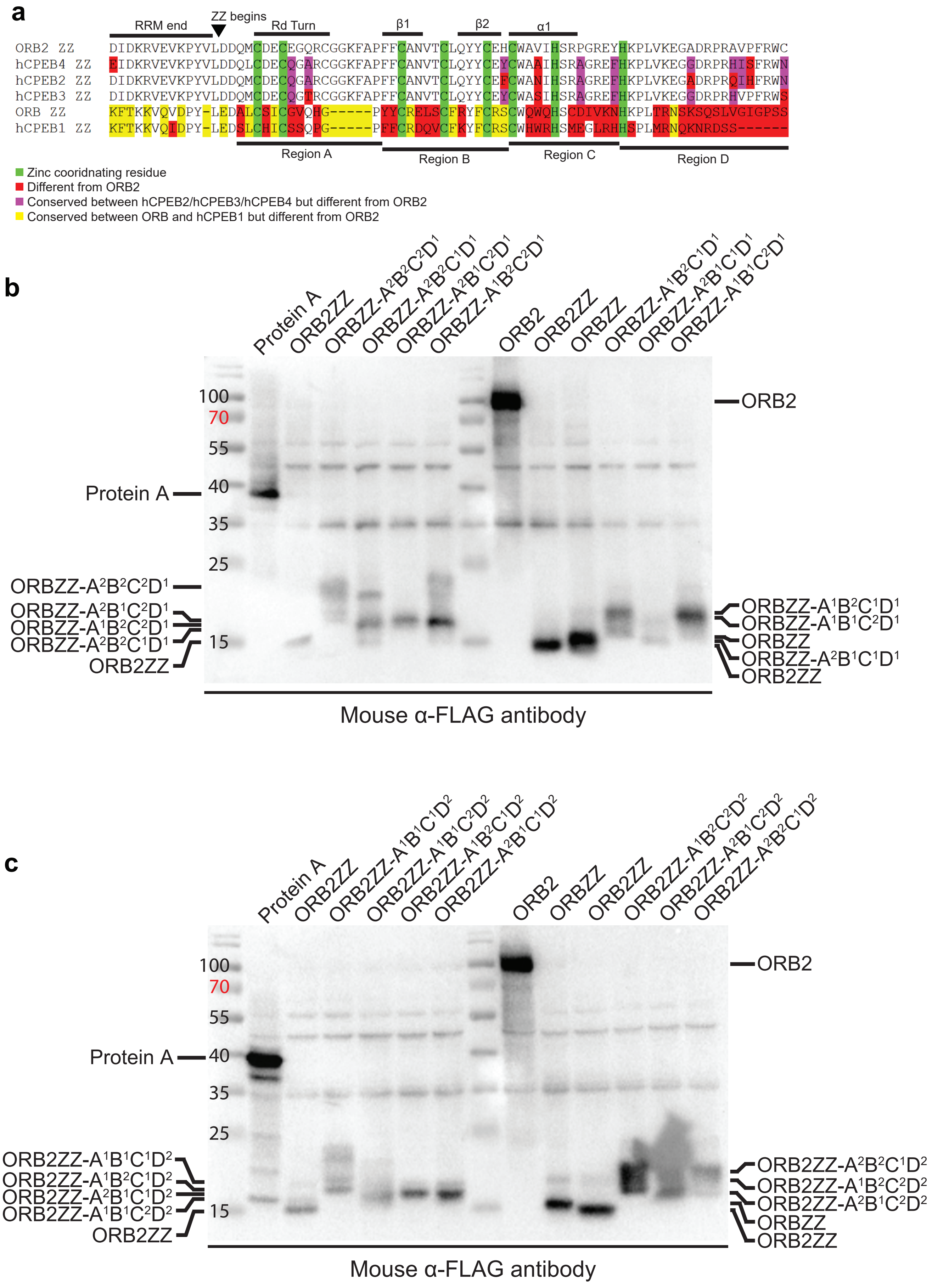

### Figure S8

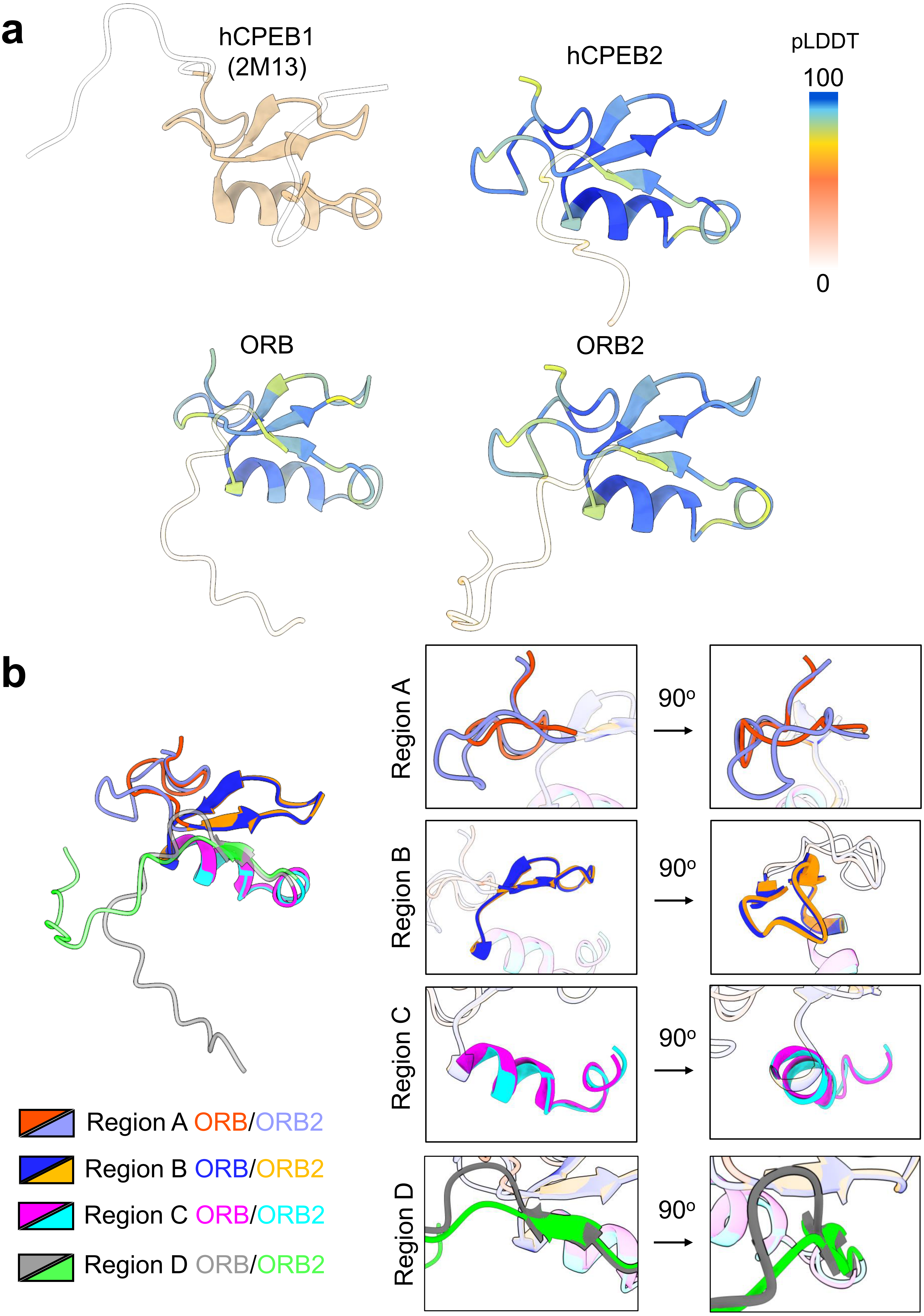

### Figure S9

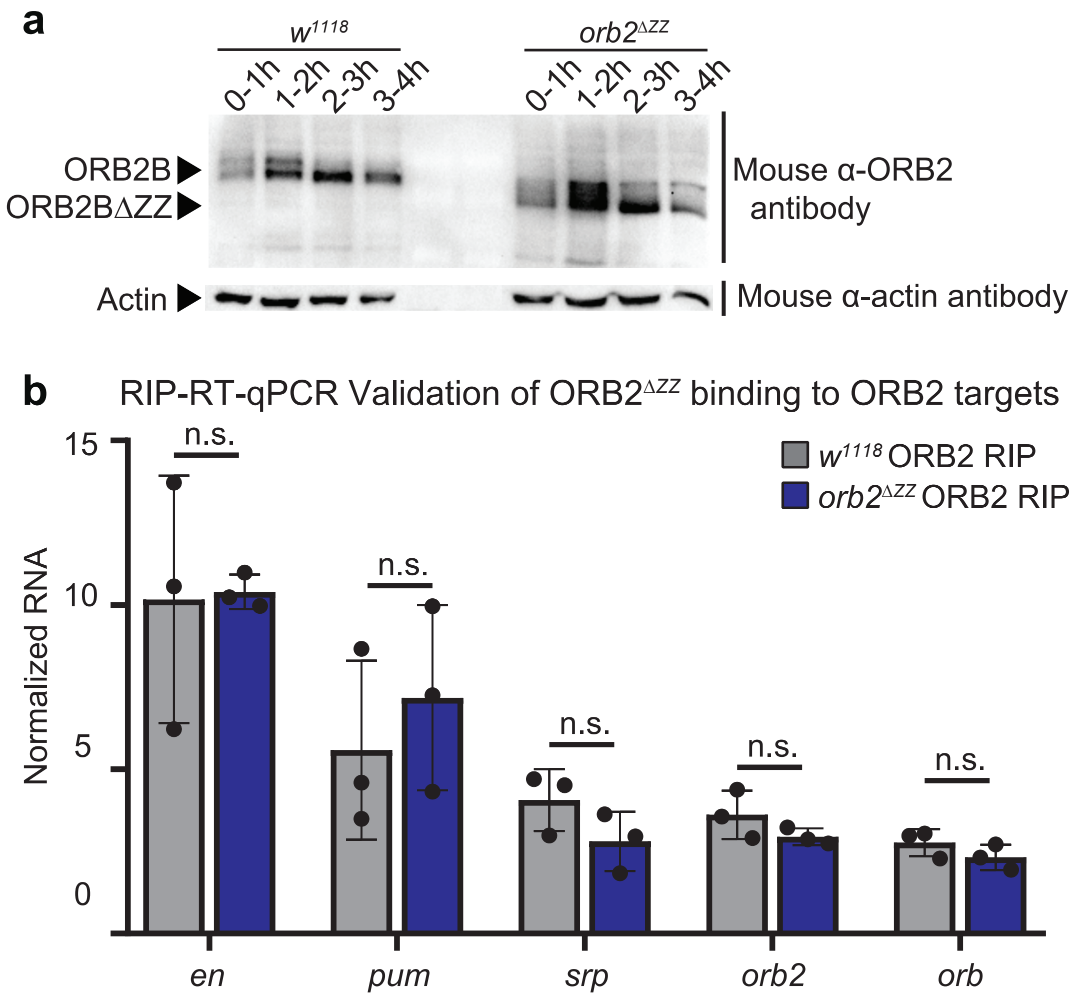

### Figure S10

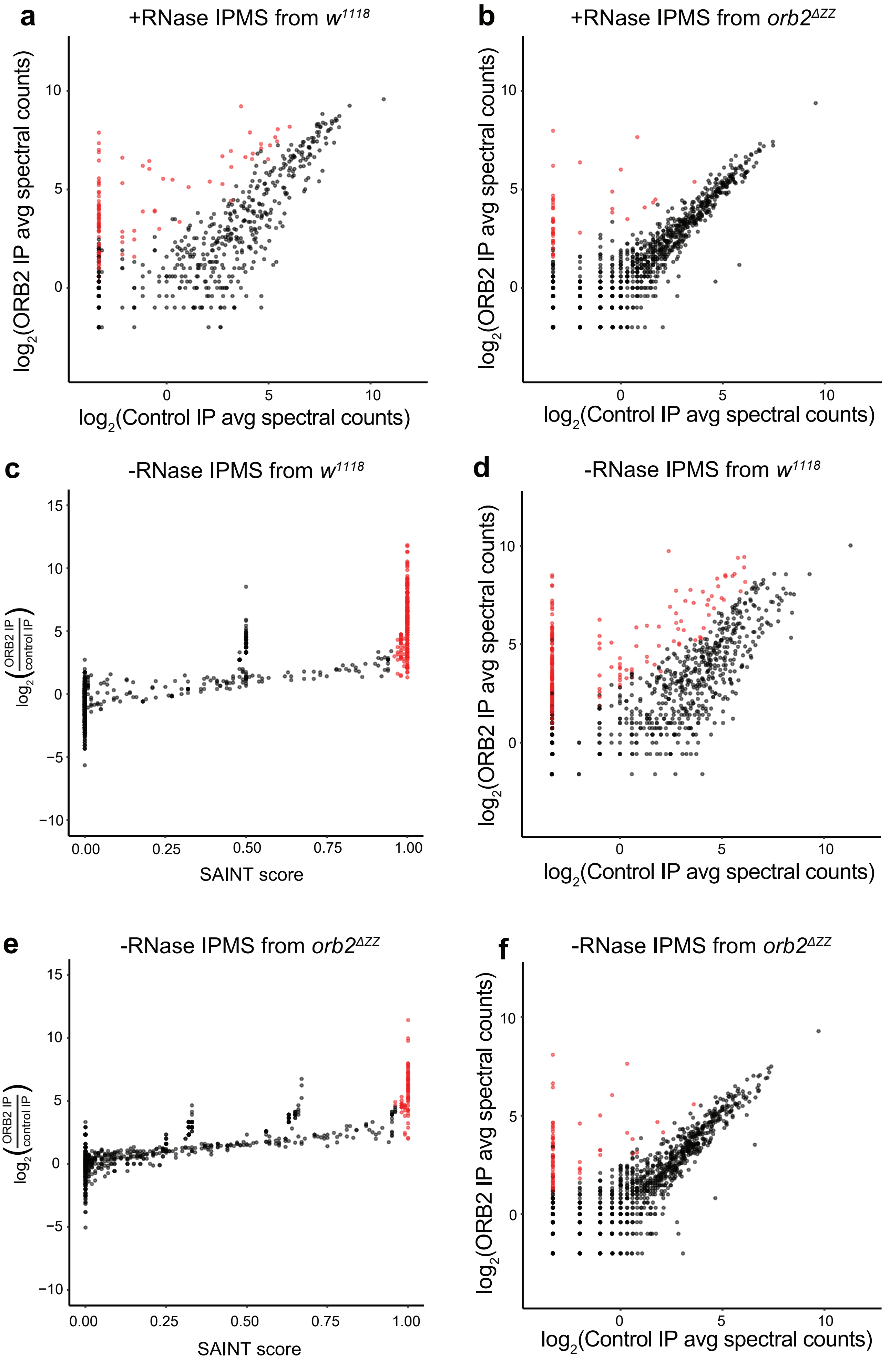

### Figure S11

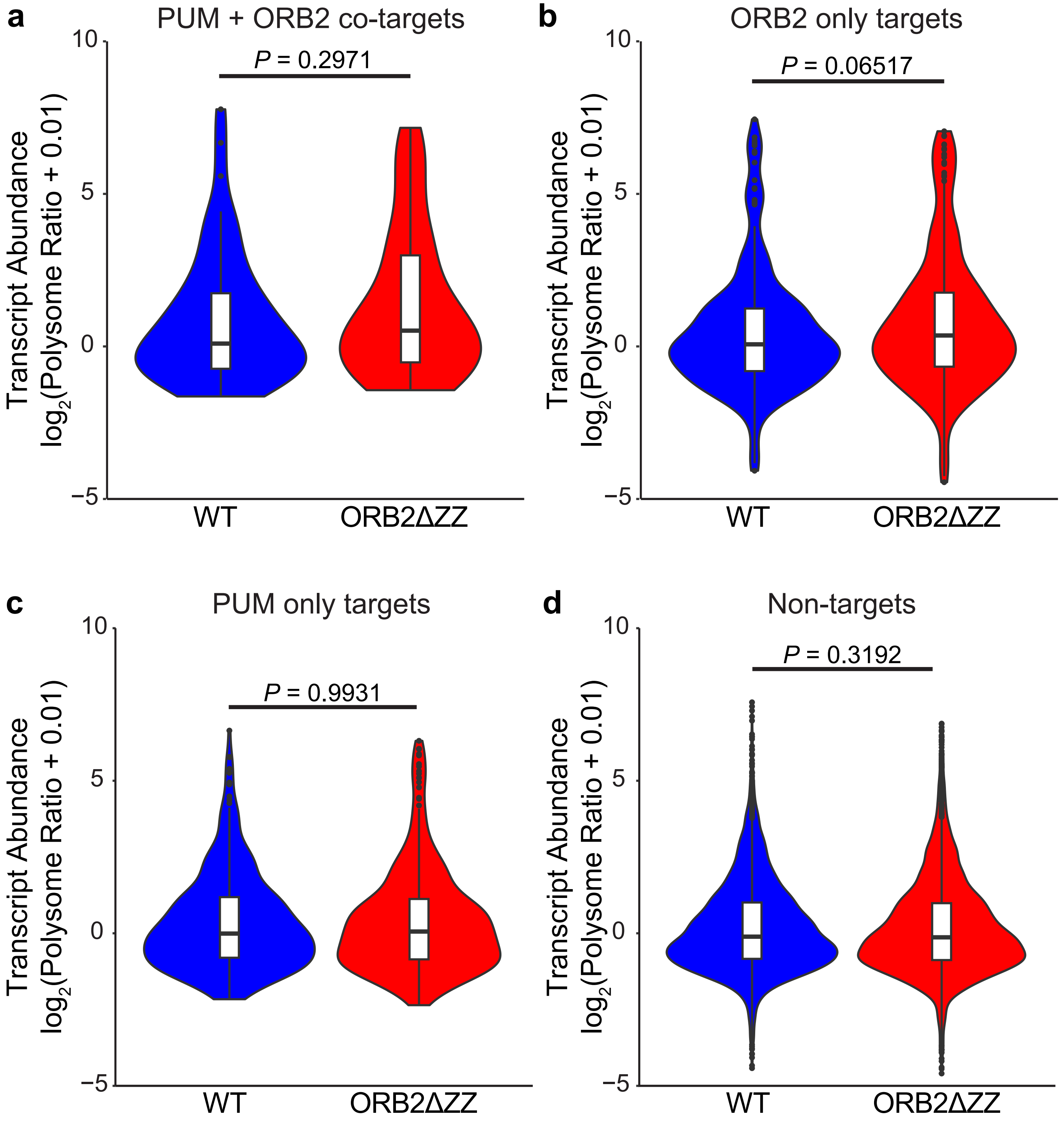

### Figure S12

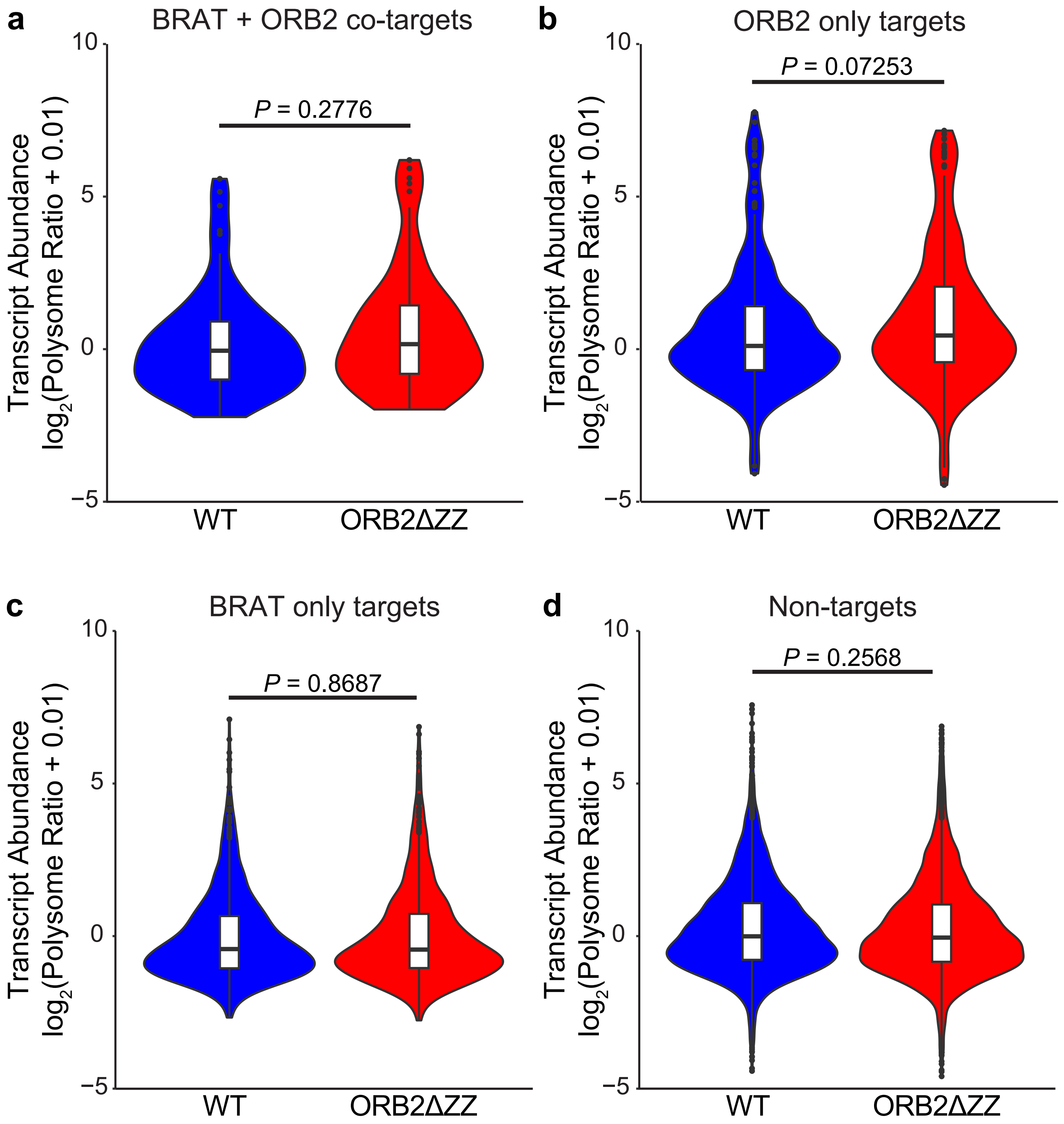
