## Supplementary material for "The ORB2 RNA-binding protein negatively regulates its target transcripts during the Drosophila maternal-to-zygotic transition via its functionally conserved Zinc-binding ‘ZZ’ domain": Figure S2

```
ORB2ΔZZ-expected     AACGGTGTTTGTGGGCGGCGTGCCACGTCCTCTGAAGGCCTTCGAACTGGCAATGATCA
                     ||||||||||||||||||||||||||||||||||||||||||||||||||||||||||||
ORB2ΔZZ-Seq          AACGGTGTTTGTGGGCGGCGTGCCACGTCCTCTGAAGGCCTTCGAACTGGCAATGATCA

ORB2ΔZZ-expected     TGGATAGATTGTACGGTGGAGTATGCTATGCTGGAATTGACACCGATCCGGAATTAAAG
                     ||||||||||||||||||||||||||||||||||||||||||||||||||||||||||||
ORB2ΔZZ-Seq          TGGATAGATTGTACGGTGGAGTATGCTATGCTGGAATTGACACCGATCCGGAATTAAAG

ORB2ΔZZ-expected     TATCCAAAGGGCGCTGGACGTGTGGCCTTCTCGAATCAGCAGAGCTACATAGCGGCCAT
                     ||||||||||||||||||||||||||||||||||||||||||||||||||||||||||||
ORB2ΔZZ-Seq          TATCCAAAGGGCGCTGGACGTGTGGCCTTCTCGAATCAGCAGAGCTACATAGCGGCCAT

ORB2ΔZZ-expected     CTCAGCCAGATTTGTGCAGCTGCAGCATGGCGATATAGACAAGCGGGTGGAGGTCAAGC
                     ||||||||||||||||||||||||||||||||||||||||||||||||||||||||||||
ORB2ΔZZ-Seq          CTCAGCCAGATTTGTGCAGCTGCAGCATGGCGATATAGACAAGCGGGTGGAGGTCAAGC

ORB2ΔZZ-expected     CCTATGTCCTTTAACGGCGGCGCTGGTAGGCCGCGCTGCACGACGAGAGACGCCAACAG
                     ||||||||||||||||||||||||||||||||||||||||||||||||||||||||||||
ORB2ΔZZ-Seq          CCTATGTCCTTTAACGGCGGCGCTGGTAGGCCGCGCTGCACGACGAGAGACGCCAACAG

ORB2ΔZZ-expected     AGGGAGCAACGACACCGGTTGTTA-AACGCAGCGCTGGCAGTGCCCAGAATCATAACAG
                     ||||||||||||||||||||||||||||||||||||||||||||||||||||||||||||
ORB2ΔZZ-Seq          AGGGAGCAACGACACCGGTTGTTTGAACGCAGCGCTGGCAGTGCCCAGAATCATAACAG
```

**NNN** = RRM sequence
NNN = ORB2 3′UTR
G to T silent mutation
Endogenous stop Codon

*ZZ domain is endogenously located between T and TAA; in the ORB2ΔZZ mutant, this sequence is removed
